# Unique potential of immature adult-born neurons for the remodeling of CA3 spatial maps

**DOI:** 10.1101/2022.09.14.507576

**Authors:** Matías Mugnaini, Mariela F. Trinchero, Alejandro F. Schinder, Verónica C. Piatti, Emilio Kropff

## Abstract

Mammalian hippocampal circuits undergo extensive remodeling through adult neurogenesis. While this process has been widely studied, the specific contribution of adult-born granule cells (aGCs) to spatial operations in the hippocampus remains unknown. Here we show that optogenetic activation of 4-week-old (young) aGCs in free-foraging mice produces a non-reversible reconfiguration of spatial maps in proximal CA3, while rarely evoking neural activity. Stimulation of the same neuronal cohort on subsequent days recruits CA3 neurons with increased efficacy but fails to induce further remapping. In contrast, stimulation of 8- week-old (mature) aGCs can reliably activate CA3 cells but produce no alterations in spatial maps. Our results reveal a unique role of young aGCs in remodeling CA3 representations, a potential that can be depleted and is lost with maturation. This ability could contribute to generate orthogonalized downstream codes supporting pattern separation.

## INTRODUCTION

Adult-born granule cells (aGCs) are added continuously to the dentate gyrus of the mammalian hippocampus, providing a unique source of plasticity absent in most other brain areas. In mice, development and maturation of aGCs takes about 8 weeks, at which time they are fully integrated to preexisting circuits and display electrophysiological properties of perinatally-born neurons (Bonaguidi et al., 2012; Groisman et al., 2020; Laplagne et al., 2006; Temprana et al., 2015; Zhao et al., 2008). By developmental week 4, aGCs go through a critical period in which they display enhanced levels of excitability and plasticity, hinting at a distinctive computational role within the dentate gyrus function (Kropff et al., 2015; Marín-Burgin et al., 2012; Mongiat et al., 2009; Stone et al., 2011; Treves et al., 2008).

Behavioral experiments suggest that immature aGCs are involved in pattern separation, aiding their CA3 target circuits to form orthogonal representations for experiences that share a high degree of similarity (Clelland et al., 2009; Goodrich-Hunsaker et al., 2008; Lee and Kesner, 2004; Luna et al., 2019; Nakashiba et al., 2012; Piatti et al., 2013; Scobie et al., 2009; Tronel et al., 2012; Yassa and Stark, 2011). In tasks that involve exploration, CA3 cells develop characteristic spatial maps, and pattern separation is expressed as remapping between similar environments (Hainmueller and Bartos, 2020; Knierim and Neunuebel, 2016; Leutgeb et al., 2007; Moser et al., 2008; van Dijk and Fenton, 2018). Consistently, remapping decays along the proximo-distal axis of CA3 in parallel to the relative prevalence of dentate inputs (Lee et al., 2015; Lu et al., 2015; Witter, 2007). While converging evidence points to a critical role of immature aGCs in pattern separation, the specific manner in which their distinctive properties induce enhanced discrimination in CA3 remains unknown.

## RESULTS

To study the influence of aGCs on CA3 representations, we performed optogenetic stimulation of young (4-week-old) or mature (8-week-old) cohorts while recording neuronal activity in free-foraging Ascl1^CreERT2^;CAG^floxStop-ChR2-EYFP^ adult mice (Figs. 1, A to C and S1, A and B). Thesemice expressed Channelrhodopsin (ChR2) in neural progenitor cells upon induction by Tamoxifen (Yang et al., 2015). Mice were then chronically implanted with optotrodes aimed at CA3. The level of expression of ChR2 and its efficacy for activating aGCs was similar in young and mature conditions (Figs. S2). After 5 days of familiarization to free-foraging in an open field arena, mice were exposed to multiple daily sessions during 4 days (Fig. 1C, days 0 to 3), including 2 sessions in a novel environment (day 2). Square and circular arenas were used as familiar or novel environments in a counterbalanced way. By day 0, spatial representations were similar across sessions due to previous familiarization, while circuit properties such as average firing rate, spatial information and stability showed no significant variations across conditions (Figs. S1C and S3 A and B). From day 1 onward, young or mature aGCs were repeatedly stimulated during one intermediate session (Fig. 1C, blue symbols) using 10-pulse trains (4 and 20 Hz), which reliably evoked spiking activity in some CA3 cells (Fig. 1D).

**Figure 1.**
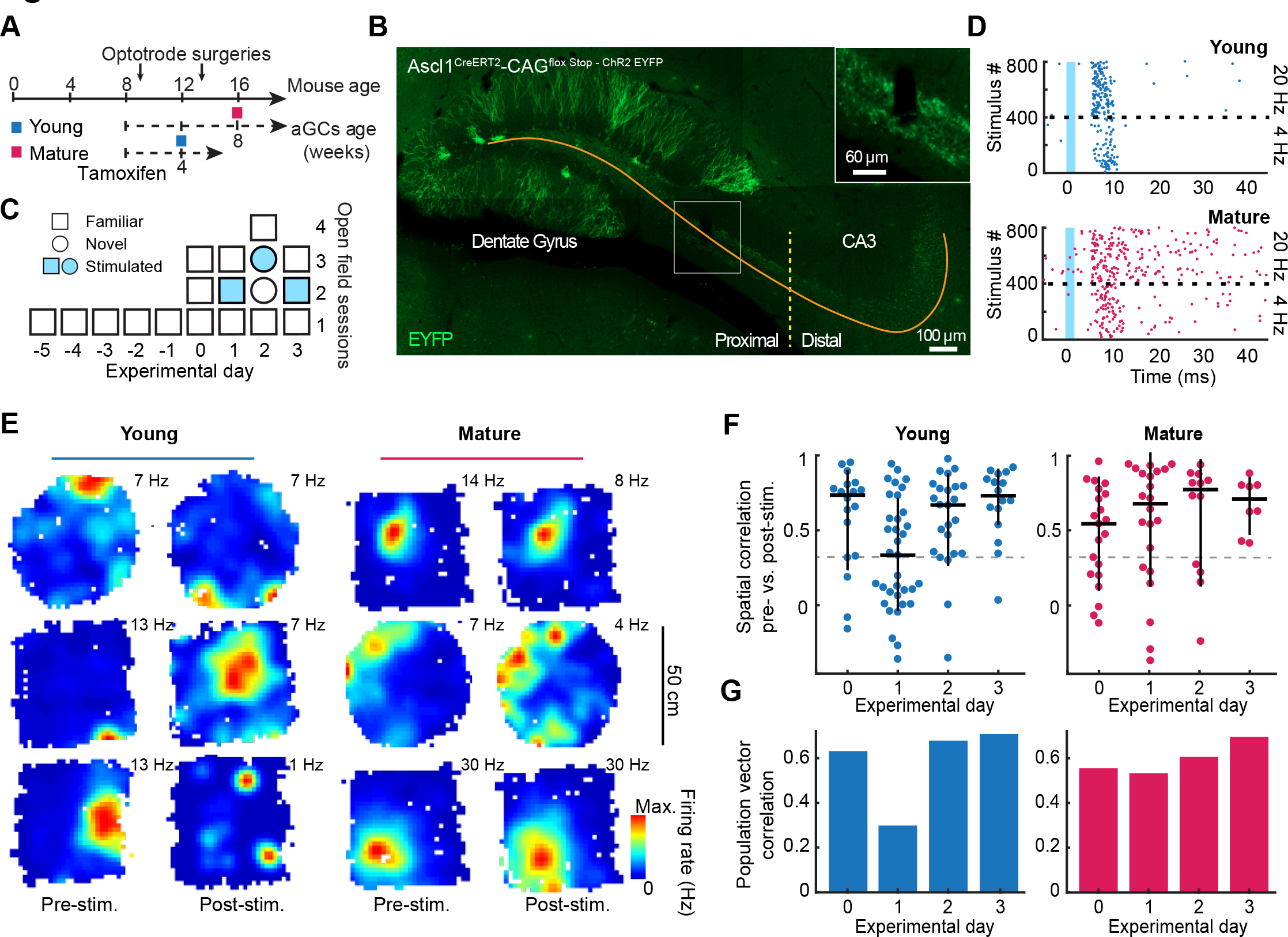
Remapping in CA3 induced by young aGC stimulation. (**A**) Schematics indicating tamoxifen injection and experimental sessions (colored squares) in mouse (top) and aGC cohort (bottom) timelines. (**B**) Representative histology showing aGCs (EYFP, green), proximo- distal axis (orange) and cut-off (yellow; 1500 µm). Inset: tetrode tip. (**C**) Behavioral sessions per experimental day in familiar or novel environments (shape code). Blue fill: stimulated session. (**D**) Raster plots showing spiking of two CA3 cells evoked by young (top) or mature (bottom) aGC stimulation. Epochs of stimulation frequency indicated. (**E**) Examples of spatial maps (color coded, maximum firing rate and session indicated) of CA3 cells (rows of subpanels) from day 1 in the young (left) and mature (right) conditions. (**F**) Distribution (one circle per neuron; black: median and IQR) of spatial correlation between pre- and post stimulation maps in the young (left) and mature (right) conditions. Cells on each day, young: 18, 35, 24 and 16; mature: 21, 22, 11 and 8. Comparisons vs. day 0 (left-tailed Mann-Whitney test) for young condition day 1: p = 0.01, other days and conditions: p > 0.25. (**G**) As (**F**) for population vector correlations. Comparisons vs. day 0 (shuffling) for young condition day 1: p = 0.04, other: p > 0.4.

To assess potential changes in spatial representation due to stimulation, we selected putative CA3 pyramidal cells that were active and stable on the first session of each day (Fig. S1D). We first analysed cells located in proximal CA3, comparing their spatial maps for the first (pre-stimulation) vs. last (post-stimulation) sessions through either spatial or population vector correlation (Fig. S1E). Remarkably, substantial remapping was observed during day 1 in several cells following stimulation of young aCGs (Figs. 1E, S1F and S4 to 6). In contrast, remapping was rare when optical stimulation targeted mature neurons, suggesting a specific role for immature aGCs in spatial processing. Across the cell population, spatial correlation was significantly lower on day 1 than on day 0 for the young condition, indicative of remapping, but similar across days for the mature condition (Fig. 1, F and G; Fig. S4 C). An increase in locomotor activity in the post-stimulation session was also observed only in the young aGC condition (Fig. S7A).

While average firing rate did not change across sessions, spiking activity transiently increased in spatial bins where young aGC stimulation had occurred, indicative of ongoing remodeling (Fig. S7B to E). In contrast to what we observed in proximal CA3, cells located in distal CA3 did not exhibit significant remapping following stimulation of young or mature aGCs (Fig. S8, A and B). For this reason, we restricted further analyses to proximal CA3 representations.

Our findings suggest that remapping in CA3 might be induced by network remodeling triggered by the activity of young aGCs. We next asked if spatial representations could be further remodeled by re-applying the stimulation protocol on subsequent days. In contrast to the observations on day 1, maps for the familiar environment remained unchanged after stimulation in a novel arena (day 2) or in the same arena (day 3; Fig. 1, F and G). These results point to a fast depletion in the capacity of young aGCs to induce remapping.

Mice in the mature condition (16 weeks old) were older than those in the young condition (12 weeks old). To rule out the possibility that remapping was related to mouse- rather than neuronal age, we stimulated a mature cohort in a group of younger mice (12 weeks old). We found no changes in spatial maps, pointing to neuronal age as the critical factor behind remapping (Fig. S9).

Our results reveal a unique potential for young aGCs to induce remapping in proximal CA3 spatial representations. This ability extinguishes once deployed and is lost with neuronal maturation.

To characterize the remodeling of CA3 spatial representations triggered by the activation of aGCs, we classified cells recorded on day 1 in the young condition into remapping and non-remapping subpopulations, according to the degree of correlation between their firing maps before and after stimulation (Figs. 1F and S1E). Using this classification, we assessed whether remapping was associated with changes in the stability of spatial maps. Stability was defined as the spatial correlation between maps constructed with data from even vs. odd segments after dividing each session into consecutive 30 s windows. Remapping and non-remapping cells displayed similar levels of stability before the stimulation session. However, while non-remapping cells continued to display high stability, remapping cells showed a marked reduction in stability after stimulation (Fig. 2, A and B; Fig. S3 C to E).

**Figure 2.**
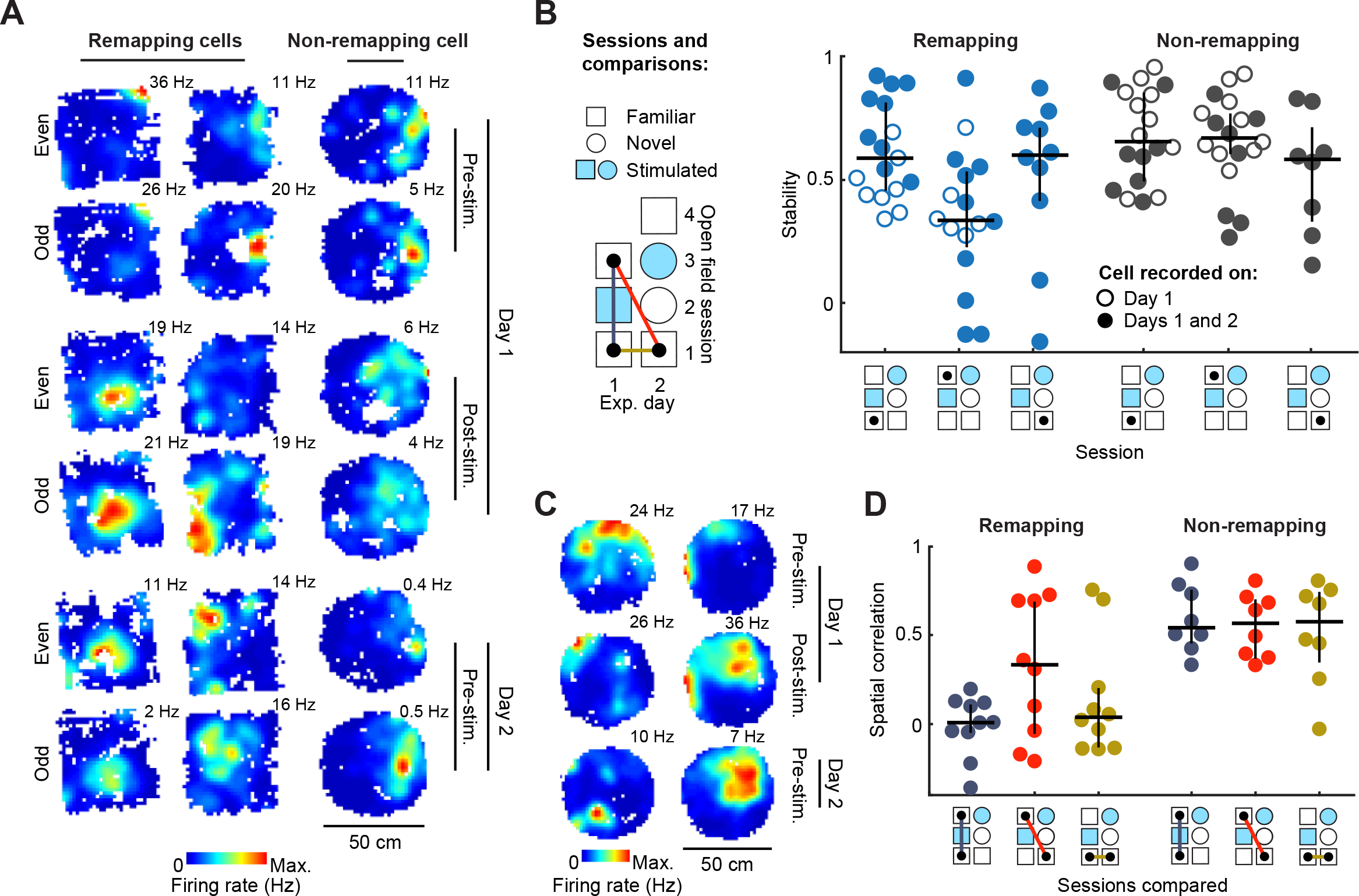
Destabilization and non-reversible remapping following the first stimulation of young aGCs. (**A**) Examples of stability for remapping (right) and non-remapping (left) CA3 cells (one per column). Spatial maps across sessions and days (color coded, maximum firing rate indicated) constructed with even (top) or odd (bottom) 30 s segments. (**B**) Left: Schematics of sessions and comparisons. Right: Distribution across sessions (indicated graphically) of stability (one circle per neuron; black: median and IQR) for remapping (blue, n=17 and non- remapping (grey; n=18) cells recorded only on day 1 (hollow) or also on day 2 (filled). Left- tailed Wilcoxon tests between columns 1 vs. 2 (all circles): p = 0.004; 1 vs. 2 (filled circles): p = 0.007; 1 vs. 3 (filled circles): p = 0.07; similar comparisons for non-remapping cells: p > 0.15. Left-tailed Mann-Whitney test between columns 1 vs. 4 (all circles): p = 0.4. (**C**) Examples of spatial maps (color coded, maximum firing rate indicated) of remapping cells (columns) across days and sessions. (**D**) Distribution of spatial correlation between maps of indicated sessions (color coded as in B; one circle per neuron; black: median and IQR) for remapping (left; n=10) and non-remapping (right; n=8) cells. Right-tailed Wilcoxon test against zero-median hypothesis for remapping cells from left to right: p = 0.46, p = 0.02, p = 0.24; for non- remapping cells p < 0.02.

To characterize the time course of this process, we monitored remapping cells on the following day and found that stability was recovered by the first session of day 2 (Fig. 2B). This recovery could indicate that remapping cells returned to pre-stimulation maps or acquired new stable maps. To distinguish between these alternatives, we analyzed the correlation between maps corresponding to different sessions of days 1 and 2 (Fig. 2, C and D). Pre-stimulation maps on day 2 were different from pre-stimulation maps on day 1, but preserved post-stimulation features. This result supports the idea that stimulation triggers the formation of new maps. Non-remapping cells, in contrast, maintained their maps throughout sessions and days.

Collectively, these results demonstrate that the stimulation of young aGCs elicits a transient loss of stability in spatial maps that is recovered within 24 h. Remapping CA3 cells ultimately form new stable maps that are uncorrelated with the original ones.

We next characterized CA3 spiking activity evoked by the same optogenetic stimulation. We visualized evoked activity of all recorded cells aligned to the stimulus onset and averaged across all stimulation pulses for young or mature aGCs (Fig. 3A, S10). Stimulation typically evoked an increase in firing rate during an early window (5 to 13 ms post stimulus) followed by a return to baseline values or brief inhibition during a late window (15 to 30 ms; Fig. S11A). These features were found in both putative pyramidal (Pyr) and fast spiking (FS) cells, defined according to their mean firing rate.

**Figure 3.**
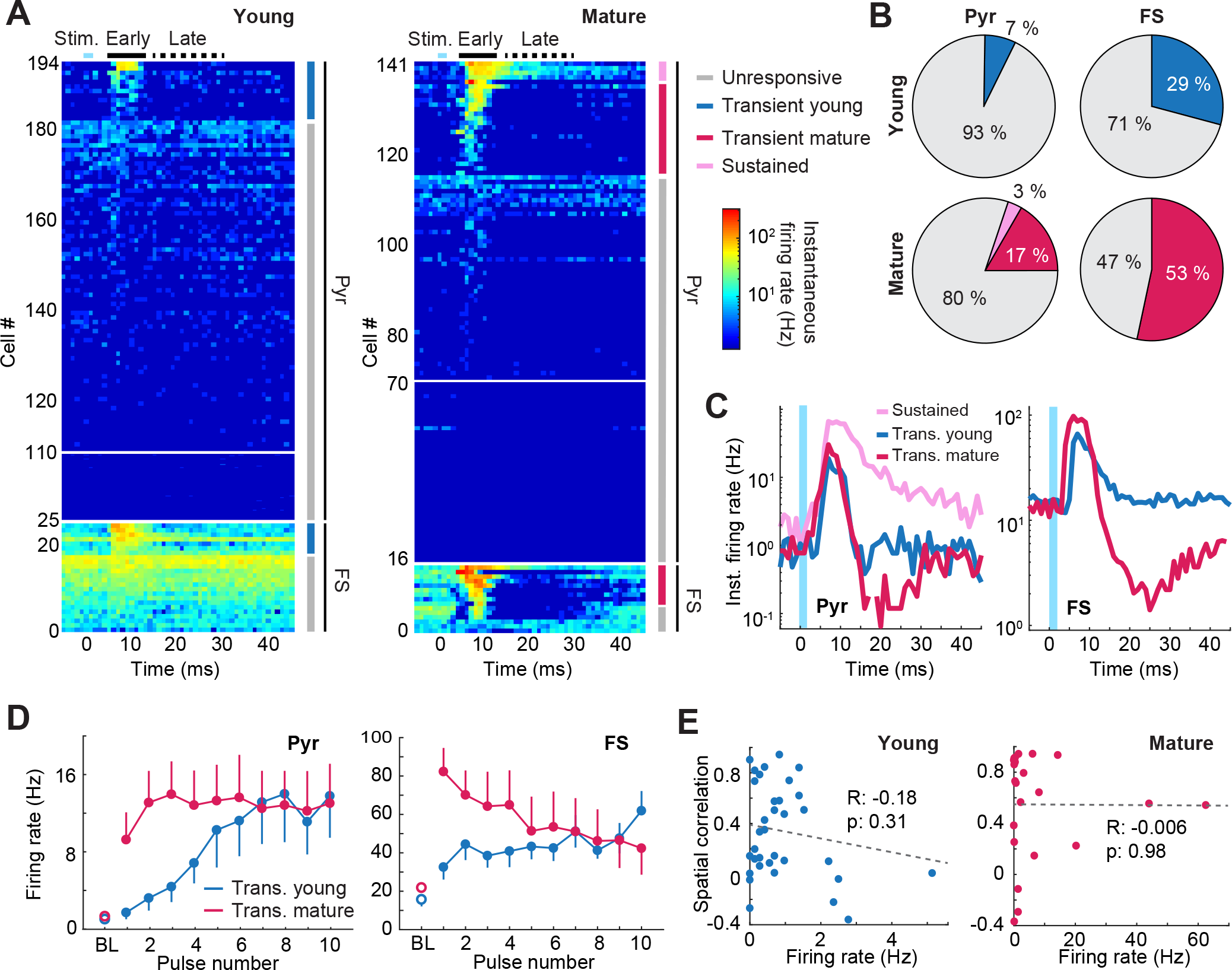
Evoked activity patterns in CA3 are not related to remapping. (**A**) Activity (color coded) relative to aGC stimulation in the young (left) or mature (right) condition for all recorded cells (rows), averaged across all stimulation pulses. Cells are ordered according to strength of response, putative cell type and response pattern (indicated). Stimulation, early and late windows are indicated. Note compression for silent cells (young: 25-109, mature: 16- 69). Pyr, transient: 13/170 (young) y 21/126 (mature); sustained: 5/126. FS, transient: 7/24 (young) y 8/15 (mature). (**B**) Distribution of response patterns (color coded as in A) for Pyr (left) and FS (right) in the young (top) and mature (bottom) conditions. Comparisons of transient cell recruitment in young vs. mature condition (shuffling): p = 0.004 (Pyr), p = 0.06 (FS). (**C**) Mean response patterns (color coded) for Pyr (left) or FS (right) cells. Comparisons of transient cell response between conditions (shuffling) for early window: p = 0.46 (Pyr), p = 0.1 (FS); for late window: p = 0.03 (Pyr), p = 0.007 (FS). Comparisons of transient (mature) vs. sustained response (shuffling): p = 0.002 (early window), p < 0.0001 (late window). (**D**) Evoked activity of transient Pyr (left) or FS (right) cells along 10-pulse stimulation trains (mean ± s.e.m; BL: baseline). Comparison of ratio between last and first pulse vs. 1 (two-tailed Wilcoxon test): Pyr p = 0.0002 (young; n=13), p = 0.02 (mature; n=21); FS p = 0.06 (young; n=7), p = 0.2 (mature; n=8). (**E**) Spatial correlation (Fig. 1F) vs. early evoked activity on day 1 in young (left) and mature (right) conditions (regression coefficients indicated).

We classified cell responses using a generalized linear model that compared activity during the early and late windows with a pre-stimulus baseline. In general, cells were classified as either “transient” (early response) or “non-responsive”. Mature aGCs recruited a higher proportion of transient cells than young aGCs, and responses were followed by a strong inhibition (Fig. 3, B and C; Fig. S8, C to E and S11, B to D). However, the amplitude of individual transient responses in both conditions was similar. In the mature condition, we found a small additional group of Pyr cells classified as “sustained”, which exhibited a remarkably strong initial response that remained elevated throughout the late window.

We also analyzed short-term plasticity of evoked spiking responses. For Pyr cells, pulse trains delivered to young aGCs elicited small initial responses with remarkable facilitation of postsynaptic spiking along subsequent pulses (about 6-fold; Figs. 3D, S8F, S11E, S12, S13). In contrast, stimulation of mature aGCs elicited a strong initial response with less facilitation (transient: 1.5-fold; sustained: 3-fold). In FS cells, responses to stimulation of young aGCs were somewhat facilitated while activity elicited by mature aGCs exhibited depression. These results reveal distinct features of synaptic transmission and plasticity during neuronal maturation and provide in vivo evidence that aGCs continue to develop their connections beyond 4 weeks of age.

We next asked if there was a relationship between the remapping observed on day 1 (Fig. 1, F to G) and the efficacy of evoked activity. Responses elicited by young aGC stimulation were low and similar for remapping and non-remapping cells (Fig. S11F). Furthermore, we found no correlation between evoked activity and remapping across the Pyr population (Fig. 3E and S11G). While remapping was not predicted by the behavior of individual CA3 cells, it was associated to prominent modifications at the network level. Compared to day 1, overall activity evoked by young aGCs changed substantially on subsequent days, resulting in a several-fold increase in the recruitment of Pyr cells (Fig. 4 and S8, G and H). In contrast, the proportion of recruited Pyr cells in the mature condition was initially high and displayed no significant changes across days. The large increase in postsynaptic responsiveness in the young condition suggests that substantial synaptic and network remodeling took place following the first stimulation of aGCs.

**Figure 4.**
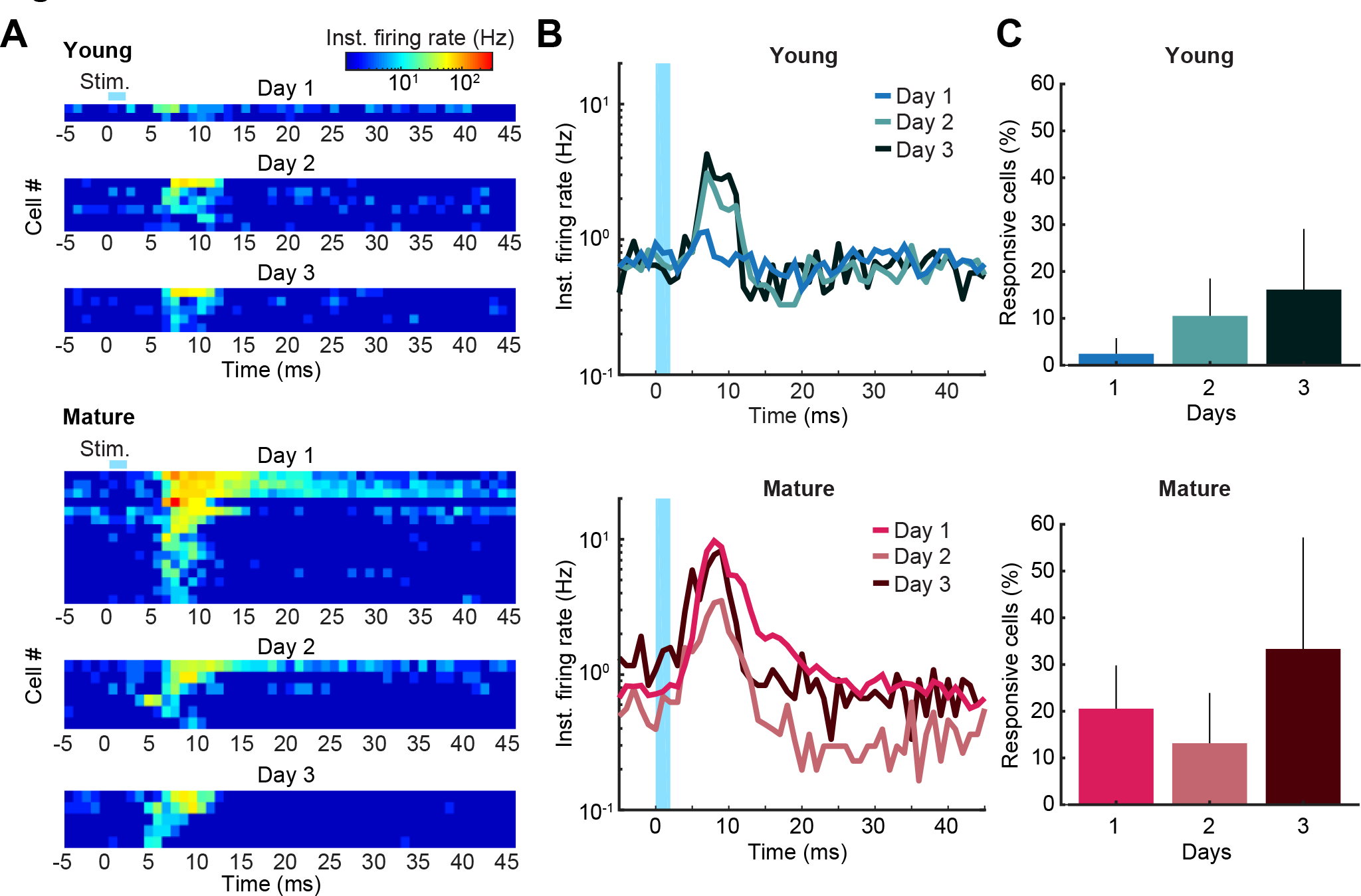
Sharp increase in CA3 recruitment following the first stimulation of young aGCs. (**A**) Evoked activity heat maps for all responsive Pyr cells (rows) across days (subpanels) in the young (top, 13 cells) or mature (bottom, 26 cells) conditions. (**B**) Evoked activity averaged over all recorded Pyr cells across days (color coded) in the young (top; n=82, 57 & 31) and mature (bottom; n=73, 38 & 15) conditions. Comparisons of early response in the young condition (shuffling): p = 0.04 (days 1 vs. 2), p = 0.03 (days 1 vs. 3). Similar comparisons in the mature condition: p > 0.1. (**C**) Percentage of responsive Pyr across days in the young (top; n=2/82, 6/57 & 5/31) and mature (bottom; n=15/73, 6/38 & 5/15) conditions. Kruskal-Wallis tests: young p = 0.03, mature p = 0.25. Dunn’s post hoc tests for the young condition: p = 0.08 (days 1 vs. 2), p = 0.02 (days 1 vs. 3).

## DISCUSSION

Our results demonstrate that stimulation of a small number of inputs from young aGCs triggers extensive network modifications. These changes are initially visualized at the level of spatial code as a destabilization and re-organization of CA3 maps. Within 24 h, postsynaptic recruitment increases several fold and cells form new stable representations of the environment, while aGCs lose their potential to induce further changes. Remapping is an indication that young aGCs trigger activity-dependent remodeling of the postsynaptic circuits, a process that might be enabled by the enhanced excitability and synaptic plasticity displayed by young aGCs (Ge et al., 2007; Gu et al., 2012; Marín-Burgin et al., 2012; Schmidt-Hieber et al., 2004). In contrast, stimulation of mature aGCs does not produce remapping, consistent with previous observations (Lee et al., 2019).

The dynamic role played by young aGCs potentially explains why remapping in CA3 may not require massive differences in population coding one synapse upstream (GoodSmith et al., 2017; GoodSmith et al., 2019; Hainmueller and Bartos, 2018, 2020; Kanter et al., 2017; Knierim and Neunuebel, 2016; Leutgeb et al., 2007; Miao et al., 2015; Schlesiger et al., 2018; Senzai and Buzsáki, 2017). Instead, activation of young aGCs by specific experiences could lead to their association with new orthogonal CA3 codes, thus supporting pattern separation (Kropff et al., 2015; Luna et al., 2019; Nakashiba et al., 2012; Scobie et al., 2009; Treves et al., 2008; Tronel et al., 2012). The transient capacity for remapping would tie experience-driven demand for novel CA3 representations to the continuous supply of new neurons as an irreplaceable source of hippocampal plasticity.

## ACKNOWLEDGEMENTS

We thank to the members of the E.K. and A.F.S. labs and for insightful discussions. A.F.S., E.K, M.T. and V.C.P are investigators in the Consejo Nacional de Investigaciones Científicas y Técnicas (CONICET). M.M. was supported by a CONICET fellowship. This work was supported by grants the National Institute of Neurological Disorders and Stroke (NINDS) and Fogarty International Center (FIC) (R01NS103758) to A.F.S.; the Argentine Agency for the Promotion of Science and Technology (PICT-2016-3611) and Pew Latin American Fellows Program in the Biomedical Sciences and Bunge & Born Foundation/ Williams Foundation/ The Pew Charitable Trusts Repatriation Grants to V.C.P; the Argentine Agency for the Promotion of Science and Technology (PICT 2019-2596) and Human Frontier Science Program HFSP-RGY0072/2018 to E.K.

## SUPPLEMENTARY MATERIALS

### METHODS

#### Animals

Experimental protocols were approved by the Institutional Animal Care and Use Committee of the Leloir Institute according to the Principles for Biomedical Research involving animals of the Council for International Organizations for Medical Sciences and provisions stated in the Guide for the Care and Use of Laboratory Animals.

Twenty-seven Ascl1^CreERT2^ – CAG ^flox^ ^Stop-ChR2^ ^EYFP^ mice (12 male and 15 female) were housed at 2-4 mice per cage from weaning. Three days before tamoxifen (TAM) injection they were transferred to individual cages, where they stayed until the end of the experiments. A running wheel was included in these cages. Animals were maintained on an inverted 12-h light/12-h schedule (lights on 08:00 pm) and tested in the dark phase. Water and food were provided ad libitum throughout the experiments.

A different set of 8 Ascl1^CreERT2^ – CAG ^flox^ ^Stop-ChR2^ ^EYFP^ mice (3 male and 5 female) was used to compare aGC counts across conditions (see section Count of ChR2 expressing cells; Fig. S2, A and B). In addition, another group of 10 Ascl1^CreERT2^ – CAG ^flox^ ^Stop-ChR2^ ^EYFP^ mice (6 male and 4 female) were used in ex vivo electrophysiology recordings to compare ChR2 efficacy across conditions (see Ex vivo electrophysiology recordings section; Fig. S2, C to E).

Experimental groups

Experimental conditions were defined by the age of the aGC cohort expressing ChR2. In the young and mature conditions, the expression was induced by TAM injections when animals were 8 weeks old. Experiments in the young condition (6 male and 6 female) were performed 4 weeks later and in the mature condition (7 male and 5 female) 8 weeks later.

In a control condition (2 male and 5 female), aimed to test mature aGCs in 12-week-old animals, TAM was injected when animals were 4 weeks old and experiments were done 8 weeks later.

#### TAM induction

TAM was delivered intraperitoneally at 120 μg/g/injection through four injections in 2 consecutive days (early morning and late afternoon) to achieve indelible expression of ChR2- EYFP in aGCs (Fig. 1A; Figs. S1B and S2A). The wheel-to-TAM period was aimed to maximize the number of labelled neurons expressing ChR2-EYFP (Yang et al, 2015).

#### Opto-drives

Tetrodes (∼25-μm in diameter) were constructed from four strands of tungsten wire (99.95% tungsten, 12.5 μm in diameter, California Fine Wire Company, USA), bound together by twisting and melting their insulation with hot air (approximately 110° C). Five tetrodes were loaded and glued into a nested assembly of stainless-steel tubes and mounted into the microdrive (Axona Limited, UK), (total weight around 2.5 g). One of the tetrodes was coupled to an optic fiber (250 μm diameter) with its tip traveling 500-700 μm below the optic fibers end. The tetrodes excited the microdrive through a guide cannula in an approximately rectangular arrangement (approximate spacing 400-μm) and every tetrode could be moved independently via a drive screw (160 μm per turn). Each tetrode was cut flat and its tip was gold-plated (gold plating solution, Neuralynx, MT, USA) to reduce the 1 KHz impedance of individual wires to 0.2–0.4 MΩ (Nano Z, White Matter, WA, USA).

#### Surgery

The surgeries to chronically implant opto-drives were performed 4 days or 32 days after TAM induction. This procedure was conducted under ketamine-xylazine anesthesia (150 μg ketamine/15 μg xylazine in 10 μl saline/g). A circular opening (approximately 1 mm in diameter) was made in the skull above the right dorsal hippocampus. The center of the craniotomy was 2.4 mm lateral to the midline and 1.7 mm posterior to bregma. Next, 7 jeweler’s screws were inserted in holes drilled in the skull around the craniotomy. After the removal of the dura, the tetrode tips were lowered 1 mm below the brain surface. The craniotomy was filled with a biocompatible transparent gel Neuroseal obtained by combining equal parts of 0.5 % sodium alginate and 10 % calcium chloride, both previously dissolved in distilled water and stored separately. The skull, the screws and the base of the opto- microdrives were then covered with auto-crystal acrylic, dental cement. After surgery animals were left to recover for 3-5 days.

#### Lowering of tetrodes

During a period of 2 – 3 weeks after recovery from surgery, tetrodes were progressively lowered toward the CA3 pyramidal layer. One of the tetrodes served as a reference and was left in an electrically-quiet zone above the hippocampus. This reference tetrode was used for differential recordings. The remaining four tetrodes served as recording probes. CA1 sharp- wave-ripple complexes and soma layer activity, visualized with Cheetah (Neuralynx, Inc., MT, USA), were the main guides to position the electrodes on the CA3 pyramidal cell layer.

#### Behavior

Free-foraging experiments were conducted in two open field (OF) arenas: a 46 cm side square and a 50 cm diameter circle (Fig. 1A and Fig. S9). Arena walls were made out of acrylic (circle) or contact paper covered cardboard (square) and acrylic floors. A white (square) or orange (circle) card (20 x 25 cm) was placed as a local cue in the south wall to facilitate orientation. Since no curtains were used, objects in the room provided additional distal cues. In sessions 2 and 3 of day 2, a novel OF was positioned in the same experimental room approximately 1.5 m away from the position of the familiar environment and the local cue card was placed in the east wall. Square and circle arenas were used as familiar or novel environments in a counterbalanced way.

A group of 4 mice (3 males and 1 female) belonging to the young condition were not sacrificed immediately after the last day of experimentation. Instead, they were kept in their home cage for around 3 weeks to repeat the battery of experiments with the same aGC cohort, this time in the mature condition. In this second round of experiments, animals were already familiarized with the square and circle environments. Thus, while the familiar environment was the same as in the first round of experiments, the novel environment was a 45 cm side equilateral triangle with contact paper cardboard walls and acrylic floor. Differences between data collected during this second round of experiments and the one collected from animals in the regular mature condition (aGCs stimulated for the first time at 8 weeks old; 4 male and 4 female) were assessed. Despite the fact that these 5 mice had additional experience with experimenter manipulations and free-foraging in the OF, and that aGCs had received more stimulation sessions, we found no differences between them and mice in the regular mature condition. Critically, we assessed remapping in both groups.

Spatial correlation between the first and last session was similar in days 1 and 0 in these 5 mice (left-tail Mann-Whitney days 1 vs. 0, p = 0.62). Similarly, there was no significant effect of mature aGC stimulation in animals belonging to the regular mature condition (left-tail Mann-Whitney days 1 vs 0, p = 0.39). Given the similarities, data from the second round of experiments of these 5 mice was pooled into the mature condition.

During all OF sessions mice were filmed from a CCTV camera (sampling rate: 25 Hz) placed on the room ceiling. Animal position was tracked from green and red LEDs attached to the headstage (HS-18-MM-LED, Neuralynx, Inc., MT, USA). In all OF sessions chocolate cereal was sporadically scattered into the arena at random locations to encourage foraging. After every OF session, the surfaces of the arena were cleaned with 70% ethanol.

Familiarization (days -5 to -1): mice were familiarized with a single 15 min presentation to the same OF arena during 5 consecutive days.

Familiar environment experiments (days 0, 1 and 3): One, two and four days after the last familiarization day, multiple presentations of the same OF took place while CA3 single unit activity was recorded. Recording sessions included: a) 10 min rest in a pedestal outside the camera range, b) three consecutive OF sessions (15 min each) separated by 5 min resting periods in the pedestal and c) 10 min rest in the pedestal. In days 1 and 3, aGCs were optogenetically stimulated on session 2 in two epochs of 4 and 20 Hz (see below).

Novel environment experiment (day 2): the protocol was similar to the one for familiar environment experiments, but there were 4 sessions. Session 1 and 4 were conducted in the familiar environment, while sessions 2 and 3 were conducted in the novel environment. All OF sessions lasted 15 min and were separated by resting periods in the pedestal. The same cohort of aGCs was optogenetically stimulated on session 3 in two epochs of 4 and 20 Hz (see below).

#### Optical stimulation

Laser stimulation was performed using a 1 W, 447 nm laser (Tolket, Argentina). The light went through a SMA to ferrule rotary joint patch cable (Thorlabs, NJ, USA) and connected to the microdrive fiber optic ferrule with a quick-release interconnect (Thorlabs, NJ, USA). Laser intensity was modulated by an ISO-Flex optical isolator (AMPI, Israel) to achieve 30 mW measured before implantation surgery at the fiber-tip with a PM160T thermal sensor power meter (Thorlabs, NJ, USA).

Optical stimulation, controlled by an Arduino Uno (Arduino, MA, USA), consisted of 80 trains of 2-ms pulses (10 pulses per train) with 5 s of inter train interval. Total energy delivered per pulse was 60 µJ. The trains were divided in two epochs of 40 trains each, with a stimulation frequency of 4 and 20 Hz, respectively. The 4 and 20 Hz simulation epochs lasted approximately 300 s and 200 s, respectively. The 4 Hz epoch started after 2 min of OF exploration and there was an exploration time in between epochs of 160 s. In this way, the 20 Hz epoch ended 2 min before the end of the 15 min session.

### Data collection

Single units activity (around 40 µV threshold) was acquired at a sampling rate of 32 KHz and filtered at 600 Hz to 6000 Hz using a Digital Lynx 4SX-M system (Neuralynx, Inc., MT, USA). Differential recordings were performed with a dedicated reference electrode in the cortex. Data processing was conducted with ad hoc MATLAB (MathWorks, MA, USA) scripts, as well as all subsequent analyses.

### Spike sorting

Events corresponding to potential neural spikes were manually clustered offline. Multiple low dimensional projections were simultaneously used to identify well separated clusters: the peak-to-peak voltage in each channel, the voltage at user-defined time points of particular channels and two dimensional tSNE reductions with default parameters (MATLAB function *tsne*). Autocorrelation and cross-correlations were used as additional separation criteria.

### Spatial maps

Spatial maps showing firing rate as a function of location were obtained by spatially smoothing (1.2 bin standard deviation 2D Gaussian filter) the ratio between spike count and time spent on a 2.5 cm side square bin grid covering all the arena.

Remapping was determined as the Pearson correlation between maps corresponding to two sessions, typically pre- and post stimulation, as indicated in each experiment. Non- covered spatial pixels were excluded for computing this and the other correlations.

Stability was computed as the Pearson correlation between maps constructed with either even or odd consecutive non-overlapping 30 s bins of each session. The total session length was segmented using 30 s bins. Mouse trajectory and single-unit activity data were grouped as falling in either even or odd bins. Maps were constructed for both groups and correlated to assess stability.

For population vector analyses maps were normalized by their mean. This ensured a purely spatial assessment of remapping, as when comparing cells individually through spatial correlation. The population vector was computed by concatenating the pixel values of all normalized spatial maps and all cells of a particular condition into a large vector. A single value, indicative of population remapping, was obtained by calculating the Pearson correlation between population vectors corresponding to two different sessions. Shuffling (see below) was used to assess statistical significance. For population vector analysis per mouse the same procedure was employed but now one PV value was obtained for each animal and then Mann-Whitney tests were performed to evaluated the statistical significance.

Spatial information rate for a spatial map with mean firing rate λ and a value λ_i_ for each of its N bins was computed as:

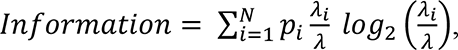

where p_i_ is the occupancy probability of bin i.

### Characterization of firing rate variation in stimulated bins

After noticing that the distribution of firing rates did not change due to stimulation in remapping or non-remapping cells, we computed for each cell the mean firing rate inside stimulated bins, normalized by the overall mean firing rate, using unsmoothed spatial maps (Fig. S7). Stimulated bins were defined as those where at least one pulse of stimulation had occurred.

### Shuffling statistics

Chance-level statistics was constructed for a given variable W (e.g. the population vector correlation) through a procedure of shuffling either the timestamps of events or the label of cells. For time shuffles, the entire sequence of spikes (Fig. S1E and D) was time-shifted by a random interval between 30 s and the session duration minus 30 s, with the end of the session wrapped to the beginning. The shuffled instance of the variable W was then calculated using the shifted events (spikes or light pulses). The collection of 100 repetitions for each cell composed the chance-level statistics. To obtain cutoff values for cell classification, shuffled data for all cells were pooled together and the 95th percentile of the pooled distribution calculated.

Label shuffles were used to assess whether a single property W was different for two groups of cells. The shuffled distribution (10000 repetitions) for W was obtained by pooling all cells and re-grouping them randomly into groups of corresponding size. In each step, a new value of shuffled W was obtained, and the actual value of W was then compared with the shuffled distribution (Figs. 1G, 3B and C, 4B; Figs. S8 D, E and G and S11 B to D and F).

### Statistical analysis

Results were expressed as the median ± and the interquartile range (IQR) or the mean ± the standard error of the mean as indicated in each caption. Data were analyzed mainly through non-parametric statistics and the different tests (Kruskal-Wallis, Friedman’s, Mann- Whitney and Wilcoxon), as well as their tails are indicated in each case. Tukey’s honest significant difference criterion was employed as post-hoc test. In all cases, p < .05 was the statistical threshold. All analyses were carried out with MATLAB (MathWorks, MA, USA) using ad hoc scripts and the available built in functions.

Regarding left-tailed test to assess remapping, the rationale is that while familiarity takes correlations between maps obtained in consecutive sessions up, remapping drives correlations down. Since our experiments asked whether aGCs can induce remapping, one- tailed tests (left-tailed in this case) are the natural statistical tool.

### Classification of patters of evoked activity

Histograms of the spiking of CA3 cells relative to single pulses of optical stimulation (-5 to 45 ms from light pulse onset) were considered as independent variables of a generalized linear model using a Poisson distribution (Figs. S9C and S11A; Fig. 3, A and B). Three categorical predictors corresponding to distinctive periods were included: baseline (-5 to 0 ms), early window (5 to 13 ms) and late window (15 to 30 ms). Significance of including early or late windows as predictors of cell activity was assessed.

### Light-pulse train analysis

Evoked activity changes along the stimulation train were computed as the ratio between last and first pulses and comparing it against the hypothesis of a median equal to 1 (two tailed Wilcoxon test; Fig. 3D). Only the early window (5 to 13 ms from light pulse onset) was considered.

### Cell selection criteria

To analyze remapping, cells that were active and stable in the first session of each day were selected. This criterion was used because remapping can only be assessed on cells with stable spatial maps. Cells were considered active if the average firing rate in this session exceed 0.1 Hz. Cells were considered stable if the stability (see above) in this session exceed the 95^th^ percentile of the shuffled distribution for that variable. Cells were considered putative fast spiking if their average firing rate during this session was above 7 Hz and putative pyramidal otherwise. Putative fast spiking cells were excluded from the analysis of remapping.

### Brain processing for histology

Electrodes were not moved after the final recording session. Subjects underwent an intraperitoneal injection of ketamine-xylazine anesthesia and were perfused intracardially with 100 ml of heparinized saline, and then with 100 ml of 4 % paraformaldehyde. Three hours after perfusion, tetrodes were raised and the brain was extracted. It was stored in 4 % paraformaldehyde overnight and then put in sucrose solution (30 %) until it precipitated (around 48 h). Frozen coronal sections (40 µm) were cut with a microtome (Leica, Germany) and dorsal hippocampal sections were mounted on glass slides. After around 20-minute drying, they were covered with PVA-DABCO mounting media and coverslips. The fluorescence microscopes BX60, Olympus (Japan; Fig. S1B) and Examiner D1, ZEISS (Germany; Fig. 1B) were used to obtain digitalized images.

### Count of ChR2 expressing cells

Immunofluorescence: Eight-week-old Ascl1^CreERT2^ – CAG ^flox^ ^Stop-ChR2^ ^EYFP^ mice were injected intraperitoneally with TAM (120 μg/g) to achieve indelible expression of ChR2-EYFP in the cell membrane of aGCs (young: n = 4; mature: n = 4). Two injections were applied on a single day instead of four injections in two days (see above) to reduce expression, which facilitated soma discrimination by reducing cell overlap. After 4 (young condition) or 8 (mature condition) weeks, mice were sacrificed. Immunostaining was done on 40 μm free- floating coronal sections throughout the brain. Antibodies were applied in TBS with 3% donkey serum and 0.25% Triton X-100. Immunofluorescence was performed using anti GFP (chicken polyclonal; 1:500; Invitrogen, MA, USA). Donkey anti-chicken Cy2 was used as secondary antibody (1:250; Jackson ImmunoResearch Laboratories, PA, USA).

Confocal microscopy: One section per animal of the septal region of the hippocampus (anteroposterior, −1.5 to −2.2 mm from bregma) was included. Images were acquired using Zeiss LSM 5 Pascal confocal microscope (Carl Zeiss, Germany). Images were acquired (20×; NA 1.3) from 40-μm-thick sections, taking z-stacks, including typically x < 0.9 μm optical slices, airy unit = 1 at 1 μm intervals. Cell counting in the default z-stack projection was restricted to cells with fluorescence intensity levels that enabled clear identification of their somata. Only EYFP^+^ cells located in the granule cell layer and sub granule zone were included in the analysis (Fig. S2, A and B).

### Ex vivo electrophysiology recordings

Slice Preparation: Eight-week-old Ascl1^CreERT2^ – CAG ^flox^ ^Stop-ChR2^ ^EYFP^ ^mice^ were intraperitoneally anesthetized with ketamine-xylazine and decapitated 4 or 8 weeks after TAM induction. To prepare transverse slices, brains were removed into a chilled solution containing (in mM): 110 choline- Cl, 2.5 KCl, 2.0 NaH_2_PO_4_, 25 NaHCO_3_, 0.5 CaCl_2_, 7 MgCl_2_, 20 dextrose, 1.3 Na^+^-ascorbate, 3.1 Na^+^-pyruvate, and 4 kynurenic acid (kyn). Coronal slices (400 μm thick) from the septal pole containing both hippocampi were cut with a vibratome and transferred to a chamber containing (in mM): 125 NaCl, 2.5 KCl, 2 NaH_2_PO_4_, 25 NaHCO_3_, 2 CaCl_2_, 1.3 MgCl_2_, 1.3 Na^+^-ascorbate, 3.1 Na^+^-pyruvate, and 10 dextrose (315 mOsm). Slices were bubbled with 95% O_2_ / 5% CO_2_ and maintained at 30°C for ∼45 min before experiments started.

*Ex-Vivo* Optogenetics: ChR2-expressing young or mature aGCs were stimulated using a 447 nm laser source delivered through the epifluorescence pathway of the upright microscope (FITC filter, 63X objective) commanded by the acquisition software. Loose-patch recordings were performed with ACSF-filled patch pipettes (8 –10 MΩ) to record optically evoked currents (Fig. 2, C to E).

## SUPPLEMENTARY FIGURE LEGENDS

**Figure S1.**
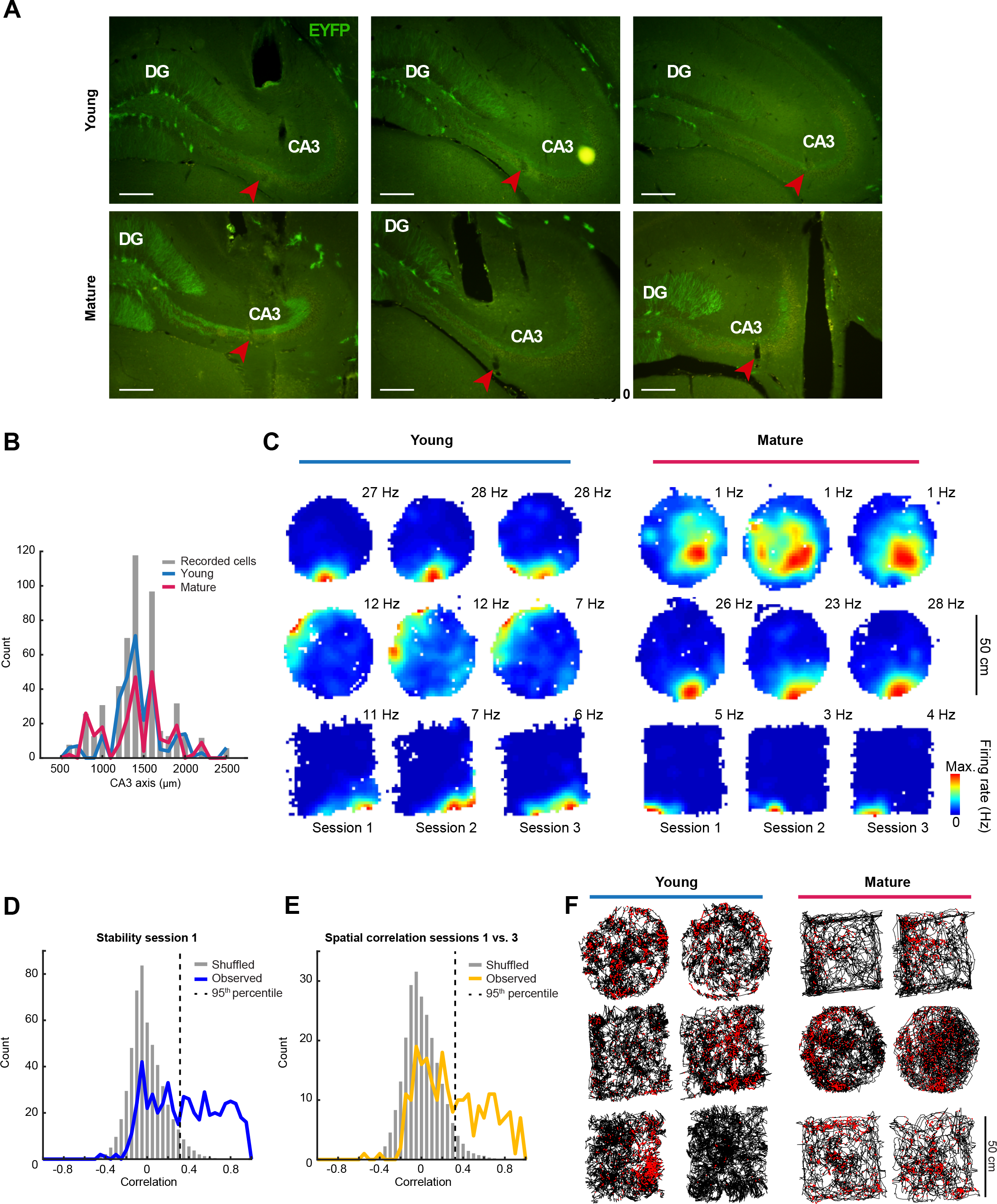
Characterization of CA3 cells and recording locations. (**A**) Representative images of coronal hippocampal histology from different implanted mice in the young (top) or mature (bottom) condition. Somata and dendritic trees of ChR2-expressing aGCs (EYFP^+^, green) can be visualized in the dentate gyrus (DG) and their mossy fibers in CA3. Red arrow: tip of tetrode near or inside the CA3 pyramidal layer. Scale: 250 µm. Top row: young aGC cohort condition. Bottom row: mature aGC cohort condition. (**B**) Distribution of all recorded cells along CA3 proximo-distal axis in the young (blue; n=293) or mature (red; n=240) aGC conditions. Grey bars: data pooled across conditions. The distributions of recording locations along the proximo-distal axis are similar for young and mature aGCs in mean (T-test p-value: 0.13) and median (Mann-Whitney p-value: 0.86). (**C**) Examples of spatial maps (color coded, maximum firing rate and session indicated) from day 0 of CA3 Pyr cells in the young (left) and mature (right) conditions (one cell per row in each subpanel). (**D**) Distribution of stability (blue line) and 100 shuffles (spike-train circular time shuffle) per neuron (grey bars; normalized by number of shuffles) during the first session of days 0 and 1 across conditions. The dashed line indicates the 95^th^ percentile of the shuffled distribution (0.32), used as a cutoff value to select stable cells. (**E**) Distribution of spatial correlation between sessions 1 vs. 3 of day 0 (orange line) and 100 shuffles (spike-train circular time shuffle) per neuron (grey bars; normalized by number of shuffles). Data pooled across conditions. The dashed line indicates the 95^th^ percentile of the shuffled distribution (0.32), used as a cutoff value to classify remapping and non-remapping cells in the young condition. (**F**) Mouse trajectory in the open field arena (black) and positions where spikes were fired (red dots). Plots depict the same cells and sessions as Figure 1E.

**Figure S2.**
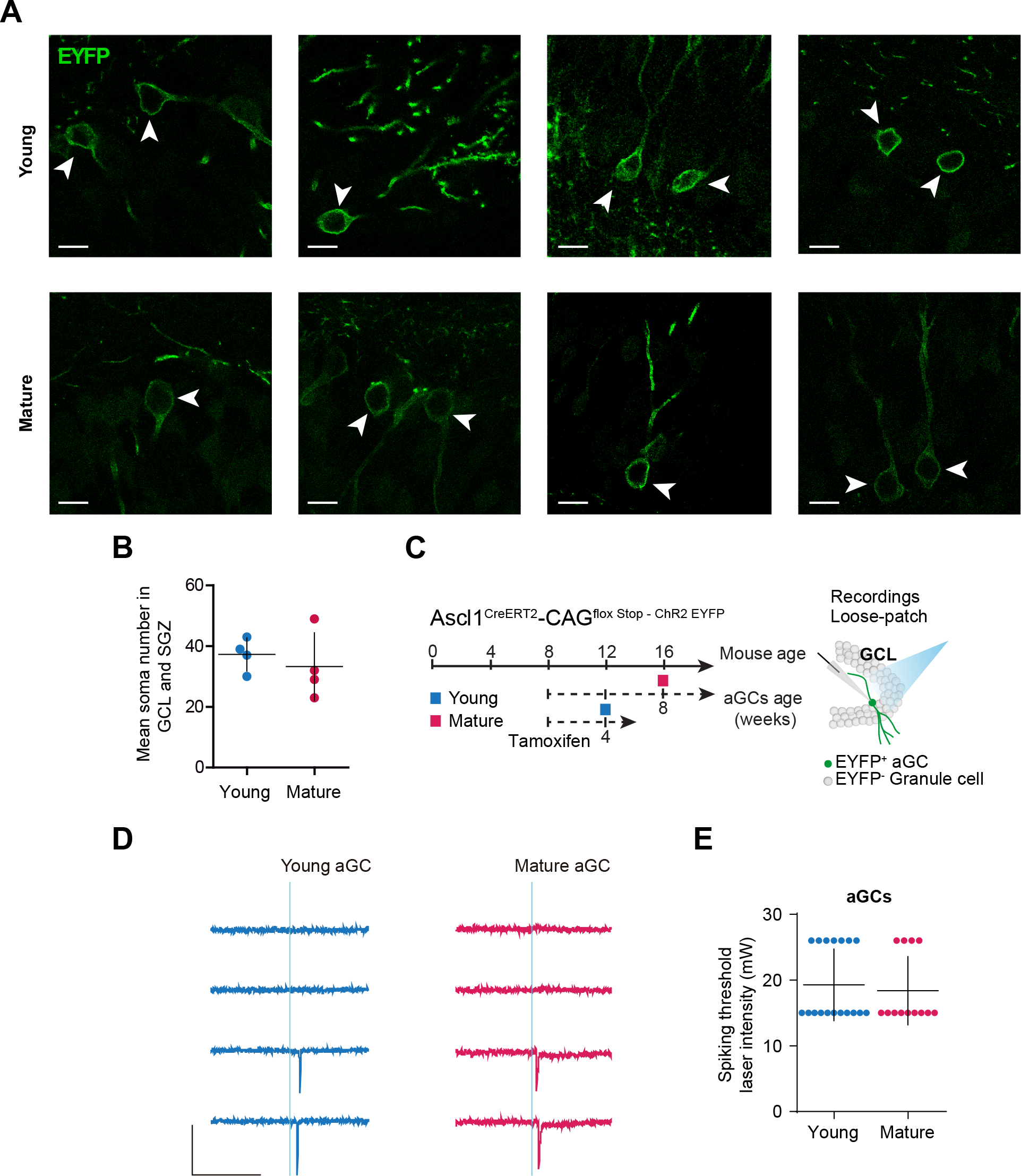
Similar expression and efficacy of ChR2 across young and mature conditions. (**A**) Representative confocal images of aGCs (arrowheads) belonging to young (top row) or mature (bottom row) cohorts. Scale: 10 µm. Each image corresponds to a different animal. (**B**) Mean soma number in the granule cell layer (GCL) and sub-granular zone (SGZ) for young (blue; mice=4) and mature (red; mice=4) aGC conditions (one mouse per circle; black: mean ± s.e.m.). Two-tailed Mann-Whitney test: p = 0.49. (**C**) Left: Experimental design similar to Figure 1A. Young adult Ascl1^CreERT2^; CAG^floxStopChR2EYFP^ mice received tamoxifen injections to induce indelible expression of ChR2 in neural progenitor cells. They were sacrificed 4 or 8 weeks later to prepare acute slices for electrophysiological characterization. Right: Schematic of loose-patch recordings on a ChR2-expressing aGC with laser stimulation. (**D**) Example loose-patch recording traces of a young (blue) and a mature (red) aGC recorded at increasing laser intensity (from top to bottom: 3, 15, 26 and 40 mW). Blue line: 2 ms stimulation. Scales: 50 pA, 100 ms. (**E**) Distribution of the spiking threshold laser intensity (mean ± s.e.m; young: 5 mice, 18 neurons; mature: 5 mice, 13 neurons). Comparison of spiking threshold across conditions: p = 0.72 (two-tailed Mann Whitney test).

**Figure S3.**
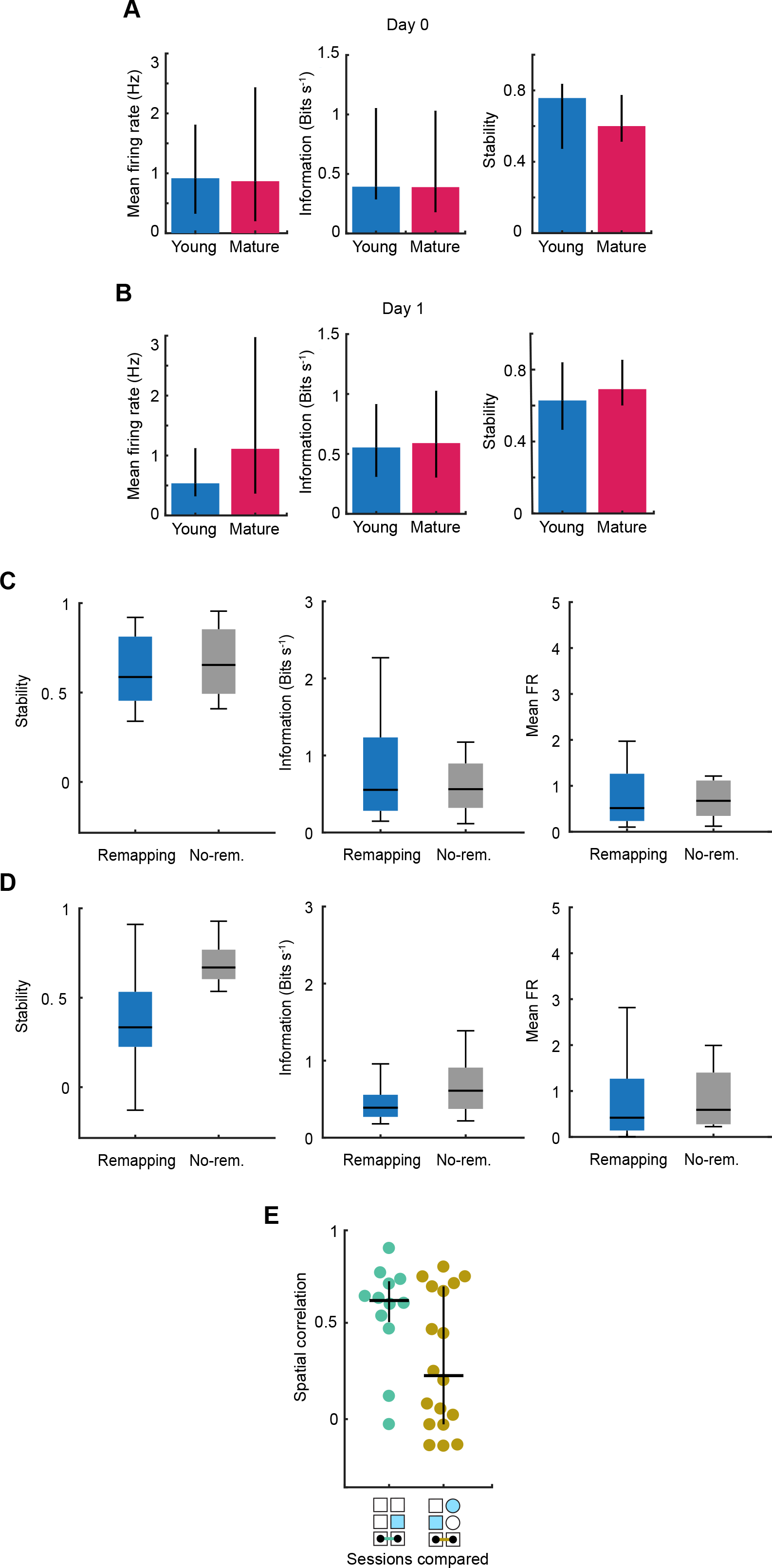
Similar pre-stimulation features for CA3 cells in the young and mature condition. (**A**) Characteristics of active and stable CA3 Pyr cells on session 1 of day 0 in the young (blue; 18 cells) and mature (red; 21 cells) conditions. From left to right: average firing rate, spatial information and stability. (**B**) Same as A but for day 1 (young: 35 cells; mature: 22 cells). No features were significantly different across conditions on days 0 or 1 (two-tailed Mann-Whitney, p > 0.11). (**C**) From left to right, boxplot (box: IQR, horizontal black line: median, lines: minimum and maximum) for stability, information and mean FR of remapping (blue) and non-remapping (grey) cells in pre-stimulation session on day 1. Two-tailed Mann-Whitney test between remapping and non-remapping cells for each property. All differences where non-significant. (**D**) As C but for cells in post-stimulation session on day 1. All differences where non-significant but post-stimulation stability (p = 0.001). Together, data in C and D point to the induced remodeling as being specific to a subpopulation of CA3 cells (i.e. typical biomarkers of global activity as mean firing rate and spatial information remained mostly unaltered). (**E**) Distribution across sessions (indicated graphically) of spatial correlation (one circle per neuron; black: median and IQR) for cells recorded in days 0 and 1 (left) or in days 1 and 2 (right). Left-tailed Mann-Whitney test: p=0.04. This result indicates that the remapping observed across days was due to the stimulation of young aGCs, rather than the consequence of a naturally occurring population drift.

**Figure S4.**
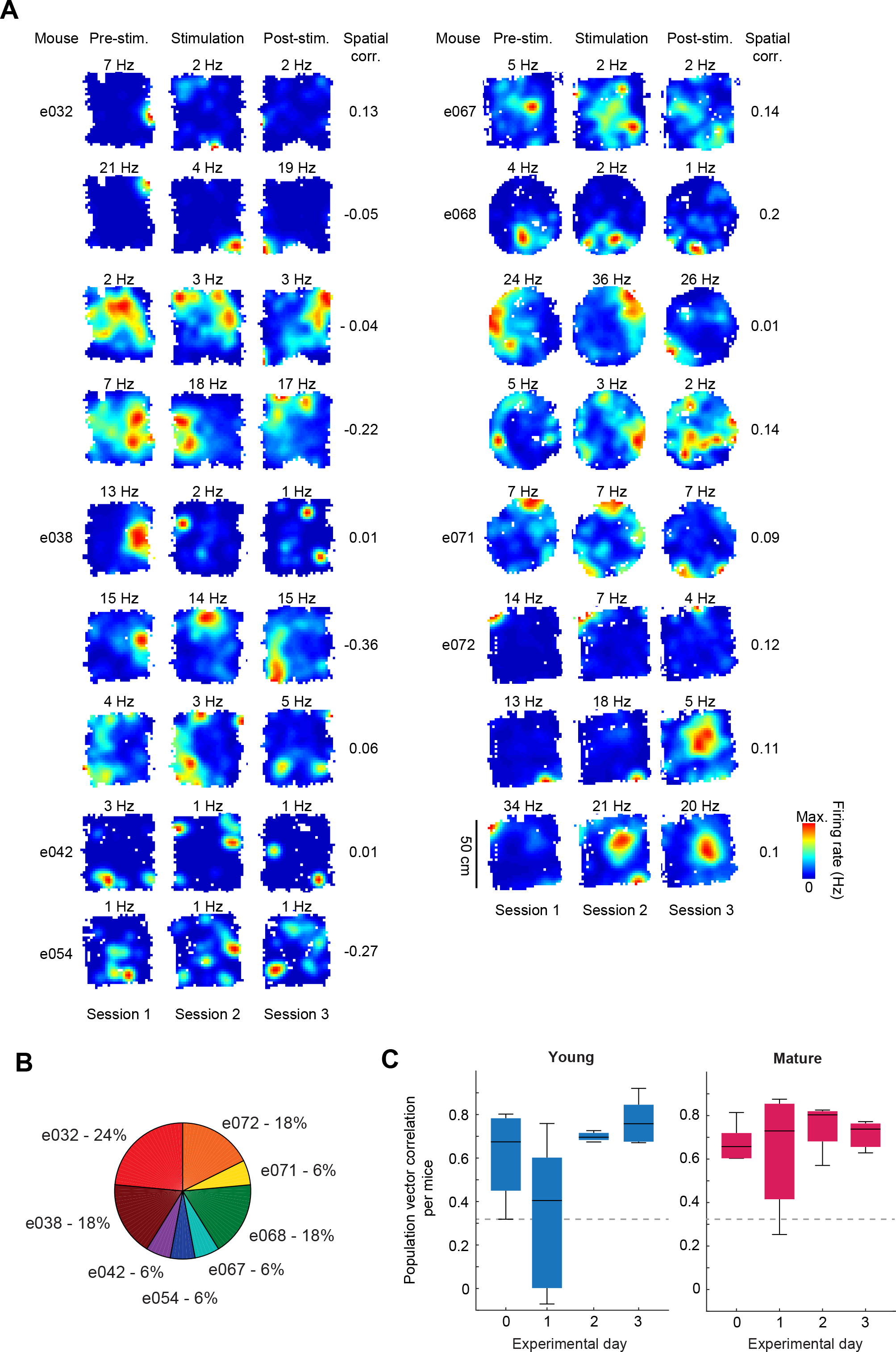
Spatial maps for all remapping cells in the young condition on day 1. (**A**) All spatial maps of CA3 active and stable Pyr cells (color coded, maximum firing rate indicated, one cell per row) recorded on day 1 and classified as remapping. Columns from left to right: mouse ID, pre-stimulation map, stimulation session map, post-stimulation map, spatial correlation between pre- and post-stimulation maps. (**B**) Distribution of remapping cells across mice. (**C)** Boxplot for the population vector correlations calculated per mouse (black: median and IQR) between pre- and post-stimulation maps in the young (left) and mature (right) conditions. Mouse number across days: 7, 11, 8 and 5 (young); 6, 9, 4 and 4 (mature). Comparisons vs. day 0 (left-tailed Mann-Whitney test) for young condition day 1: p = 0.03, other days and conditions: p > 0.36.

**Figure S5.**
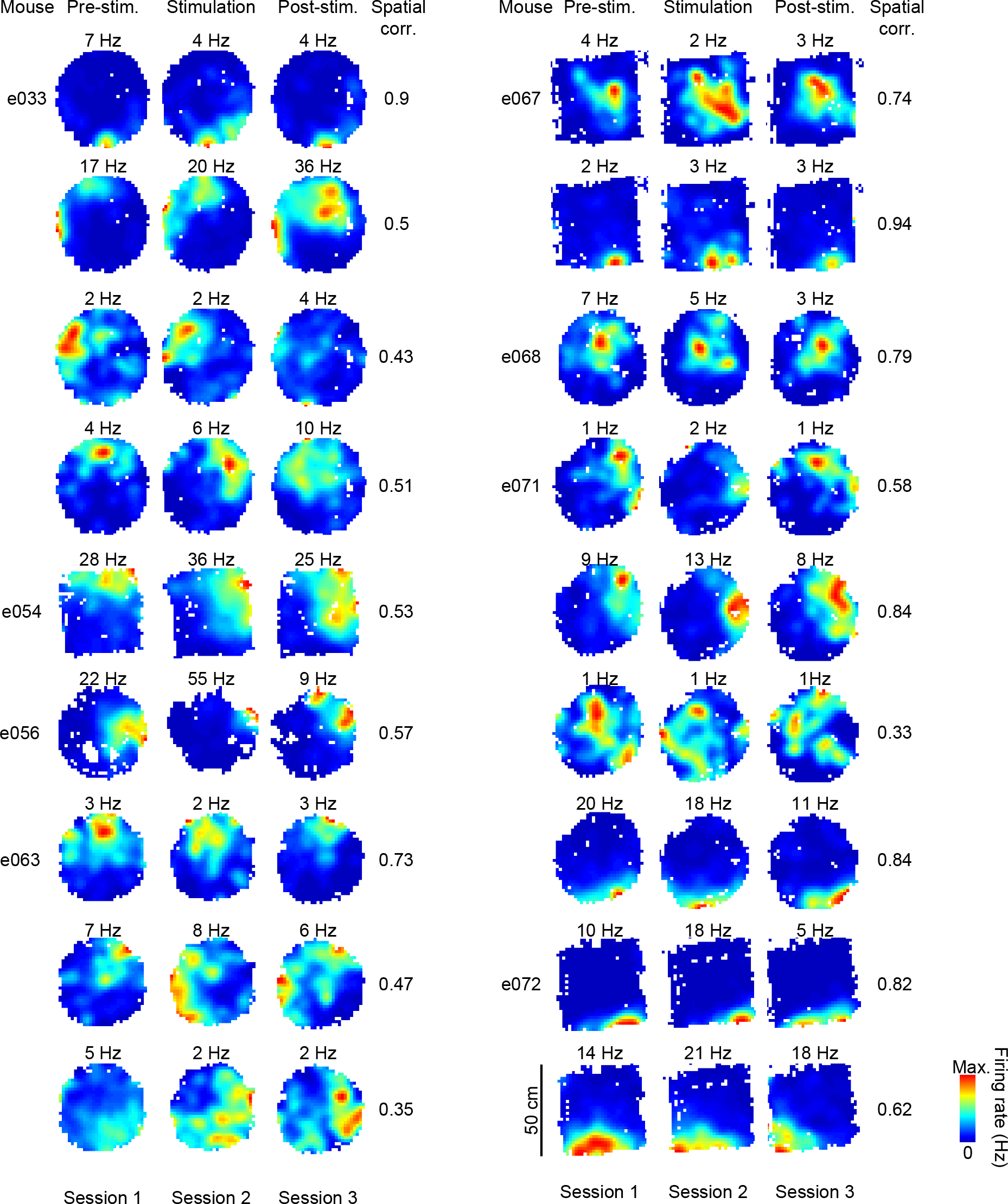
Spatial maps for all non-remapping cells in the young condition on day 1. All spatial maps of CA3 active and stable Pyr cells (color coded, maximum firing rate indicated, one cell per row) recorded on day 1 and classified as non-remapping. Columns from left to right: mouse ID, pre-stimulation map, stimulation session map, post-stimulation map, spatial correlation between pre- and post-stimulation maps.

**Figure S6.**
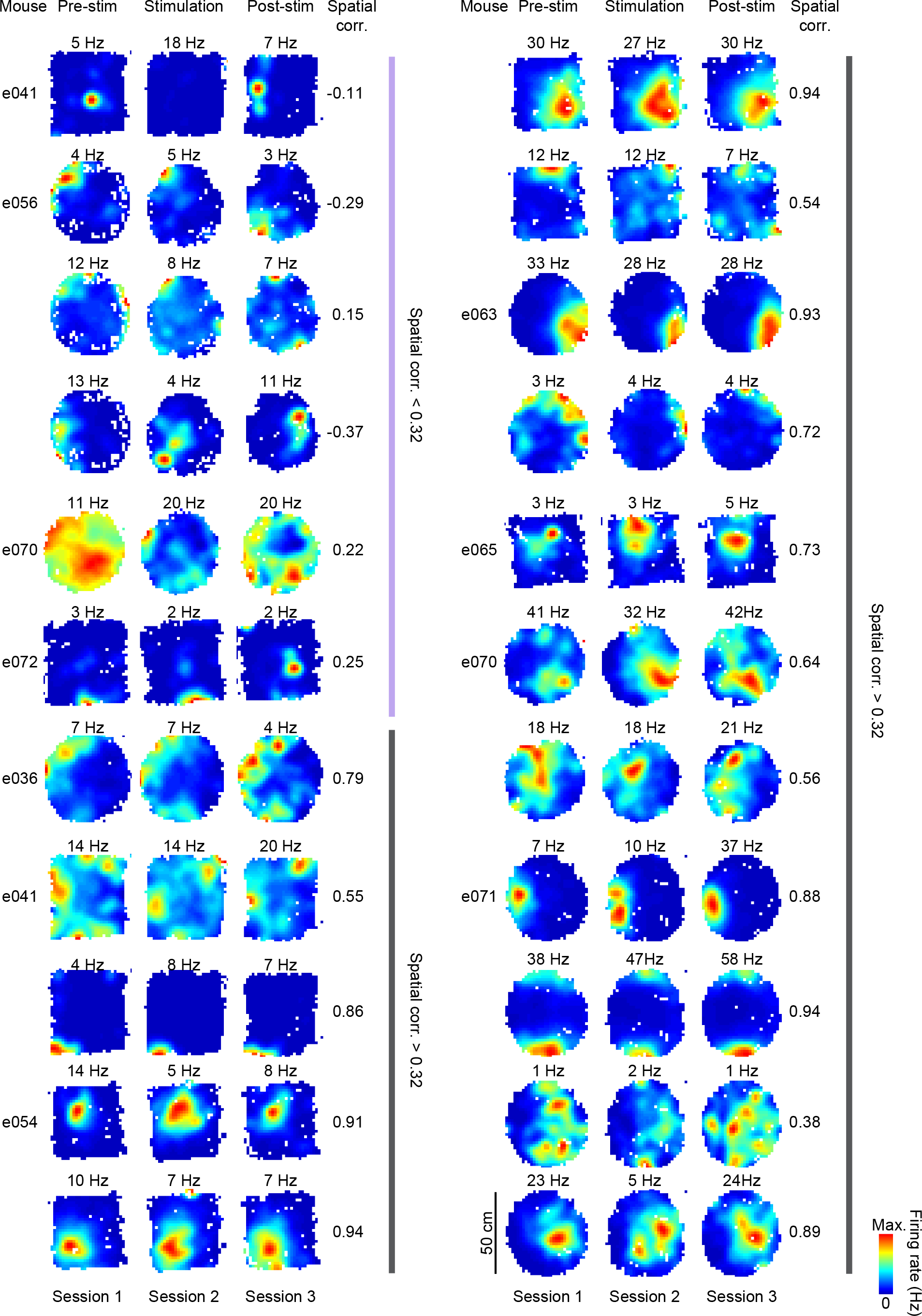
Spatial maps for all active and stable cells in the mature condition on day 1. All spatial maps of CA3 active and stable Pyr cells (color coded, maximum firing rate indicated, one cell per row) recorded on day 1. Columns from left to right: mouse ID, pre-stimulation map, stimulation session map, post-stimulation map, spatial correlation between pre- and post-stimulation maps. Cells are grouped according to whether their spatial correlation was below (purple) or above (grey) the cutoff value of 0.32.

**Figure S7.**
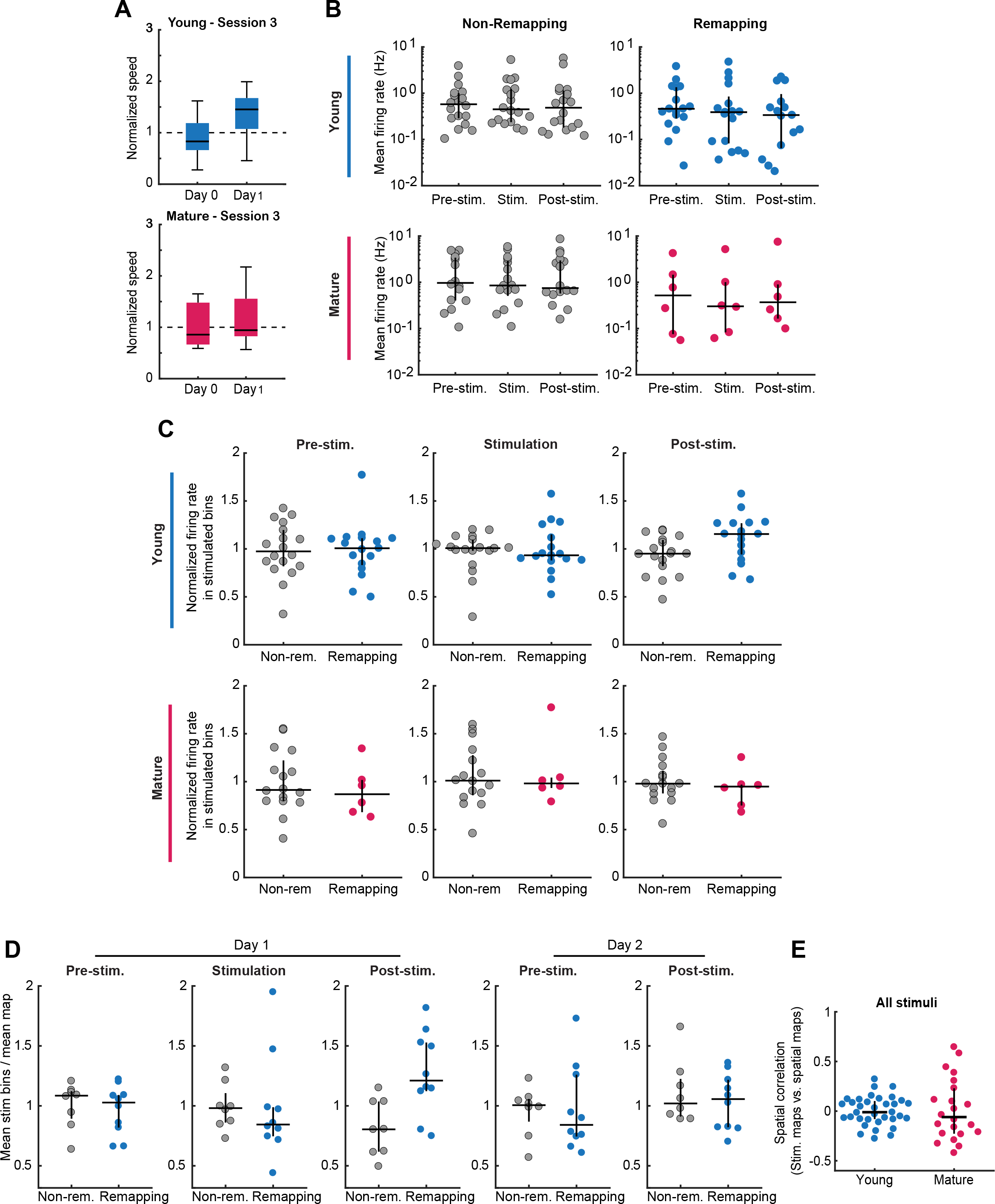
aGCs stimulation affects locomotion, firing rate and place field reorganization. (**A**) Mean normalized speed during session 3 (post-stimulation) in days 0 and 1 in the young (left) and mature (right) conditions. Comparison of normalized speed against one (two-tailed Wilcoxon test): young p = 0.5 (day 0), p = 0.02 (day 1); mature p = 0.73 (day 0), p = 0.56 (day 1). These results show an increase in locomotion after young aGCs stimulation. (B) Mean firing rate across sessions of day 1 for CA3 cells with spatial correlation above (left, non-remapping; n=18) or below (right, remapping; n=17) the classification cutoff value (0.32; Fig. S1E) in the young (top) or mature (bottom) condition. No variation across sessions was found (Kruskal-Wallis tests: p > 0.5). (**C**) Subpanels show the distribution of relative firing rate inside stimulated bins across sessions of day 1 (columns; indicated) for remapping (blue or red) and non-remapping (grey) cells in the young (top) and mature (bottom) conditions. Spatial bins that received stimulation on session 2 were identified and used to calculate relative firing rate in all sessions. For each cell and session, the normalized firing rate was calculated as the ratio between the firing rate inside stimulated bins and the mean firing rate over the whole session (one circle per cell, black: median ± IQR). Comparison between remapping and non-remapping cells in the post-stimulation session: p = 0.024; similar comparisons for other sessions or condition: p > 0.44 (two-tailed Mann Whitney test). These results show that there was a transient increase in relative spiking inside stimulated bins for remapping cells. (**D**) Same as C but including pre- and post-stimulation sessions of day 2 for cells recorded on both days 1 and 2. Only the difference between remapping (n=10) and non-remapping (n=8) cells on post-stimulation session of day 1 was statistically significant (p = 0.009). (**E**) New maps are not correlated with stimulation locations. Distribution (one circle per neuron; black: median and IQR) of spatial correlation between post stimulation spatial maps (session 3) and stimulation position maps (session 2) in the young and mature conditions in day 1. Comparisons against 0 (two-tailed Wilcoxon test) for young (All cells, p = 0.8; remapping, p = 0.64; non-remapping, p = 0.47 - last two not shown) and mature (All cells, p = 0.91; remapping, p = 1; non-remapping, p = 0.95 - last two not shown). This result indicates that, despite the transient increase in relative spiking shown in A to D, there is no a direct relation between stimulation location and spatial map reorganization.

**Figure S8.**
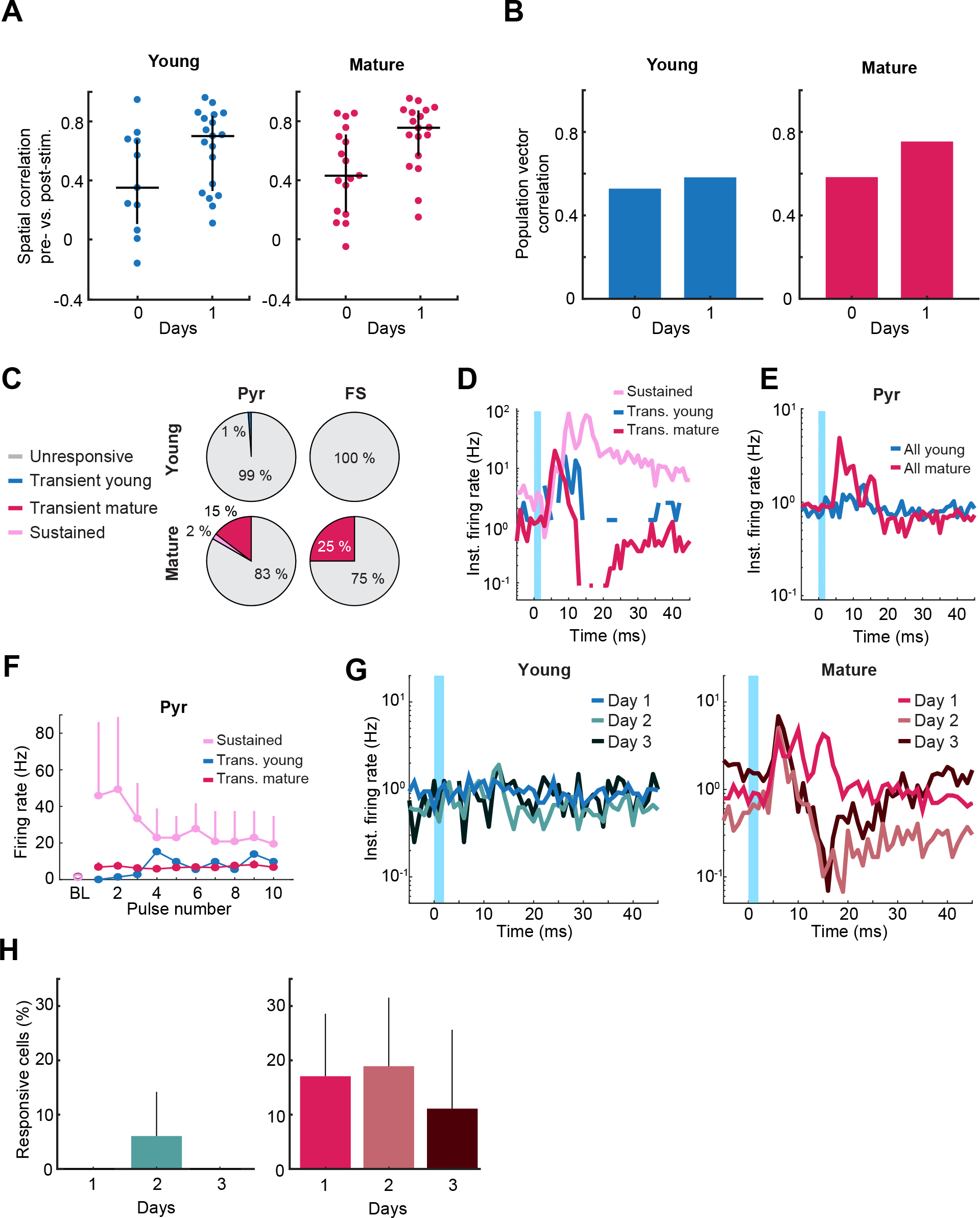
No remapping and low levels of activity evoked by young aGC stimulation in distal CA3 cells. (**A**) Distribution as in Fig. 1F of spatial correlation for all active and stable distal CA3 cells on days 0 and 1 in the young (left, blue, 11 & 19 cells) and mature (right, red, 17 & 18 cells) conditions (black: median ± IQR). Comparisons between days 0 and 1 (left-tailed Mann-Whitney test): p = 0.96 (young), p = 0.99 (mature). (**B**) Population vector correlation for the same days and cells in the young (left) and mature (right) conditions. Comparisons between days 0 and 1 (shuffling): p = 0.65 (young), p = 0.97 (mature). (**C**) Distribution of evoked response patterns in distal CA3 (color code as in A) for Pyr (left) and FS (right) in the young (top; n=1/88 & 0/10) and mature (bottom; n=14/96 & 1/4) conditions; sustained: 2/96. Comparisons of transient cell recruitment between conditions (shuffling): p = 0.0004 (Pyr), p < 0.0001 (FS). (**D**) Mean response patterns (color coded) of distal Pyr cells. Comparison of transient cell response between conditions (shuffling): p = 0.8 (early window), p = 0.8 (late window). Comparisons of transient (mature) vs. sustained response (shuffling): p = 0.009 (early window), p < 0.0001 (late window). Broken lines correspond to a value of 0, which cannot be ploted in log scale. (**E**) Mean evoked activity of all recorded distal Pyr cells in the young (blue; n=88) or mature (red; n=96) condition. Comparisons between conditions: p = 0.003 (early window), p = 0.9 (late window). (**F**) Evoked activity of transient young (blue), transient mature (red) and sustained (pink) Pyr cells along 10-pulse stimulation trains (mean ± s.e.m; BL: baseline). Comparison of ratio between last and first pulse vs. 1 (two-tailed Wilcoxon test): p = 0.09 (mature transient Pyr; n=14). Note difference with the facilitation observed in proximal CA3 for mature transient cells (Fig. 3D). Other plots and statistics were not included due to the low number of data points (young transient Pyr: n = 1; mature sustained: n = 2; mature transient FS: n = 1). (**G**) Evoked activity averaged over all recorded distal Pyr cells across days (color coded) in the young (blue; n=46, 32 & 10) or mature (red; n=41, 37 & 18) conditions. Comparisons of early response between days 1 vs. 2 and days 1 vs. 3 in both conditions (shuffling): p > 0.19. (**H**) Percentage of transient distal Pyr across days in the young (left) and mature (right) conditions. Kruskal-Wallis tests: p = 0.17 (young), p = 0.76 (mature).

**Figure S9.**
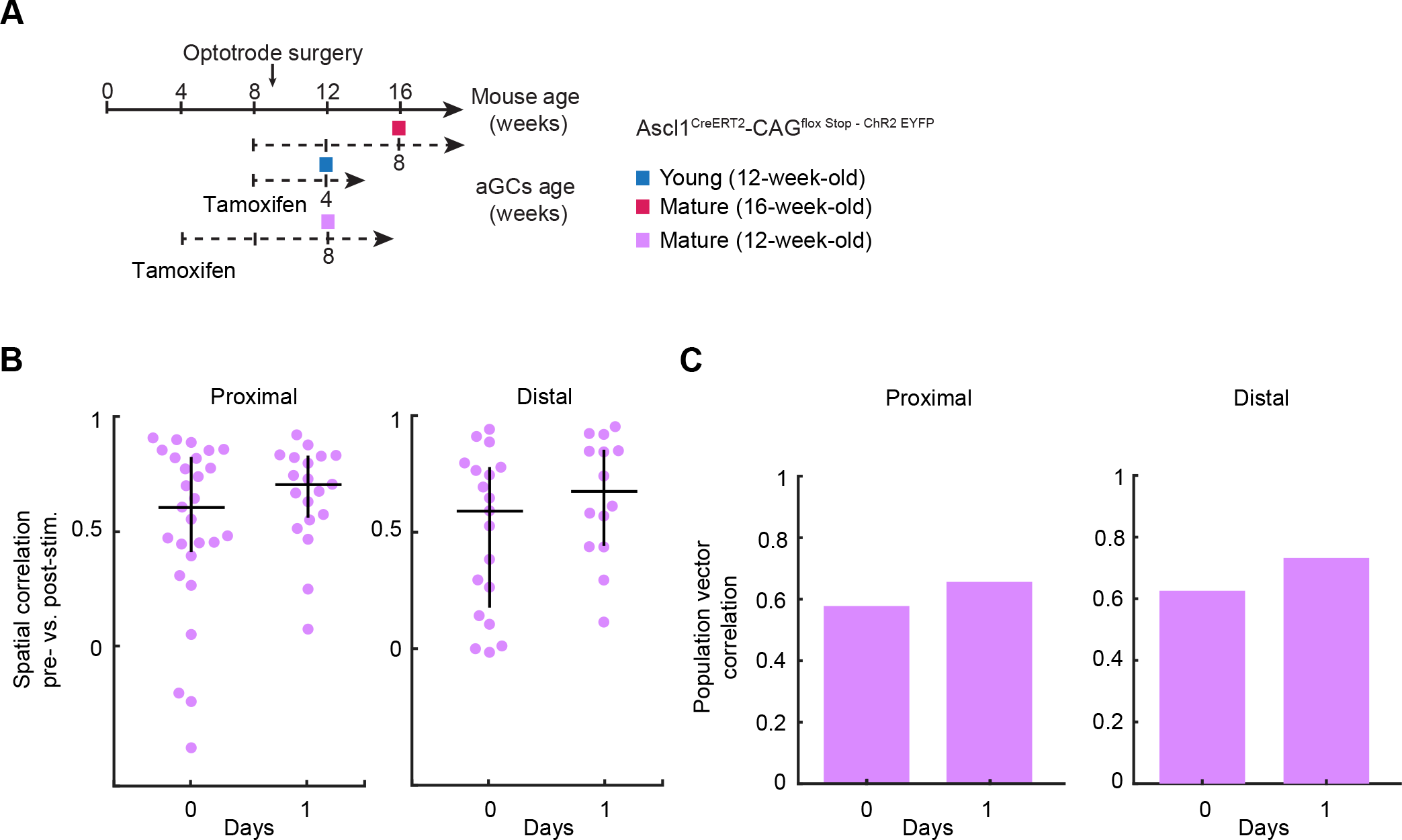
No remapping following the stimulation of a mature aGC cohort in 12-week-old mice. (**A**) Schematics as in Fig. 1 A but adding the timeline for the 12-week-old mice control (purple). Four-week-old transgenic mice received tamoxifen injections and experiments were done 8 weeks later, when animals were 12 weeks old and aGCs expressing ChR2 were mature. (**B**) Distribution of spatial correlation, as in Fig. 1F, for all active and stable cells on days 0 and 1 in proximal (left, 19 & 14 cells) and distal (right, 27 & 19 cells) CA3 (one circle per cell; black: median ± IQR). Comparisons between days (left-tailed Mann-Whitney test): p = 0.85 (proximal), p = 0.95 (distal). (**C**) Population vector correlation for the same days and cells in proximal (left) and distal (right) CA3. Comparisons between days (shuffling): p = 0.99 (proximal), p = 0.91 (distal).

**Figure S10.**
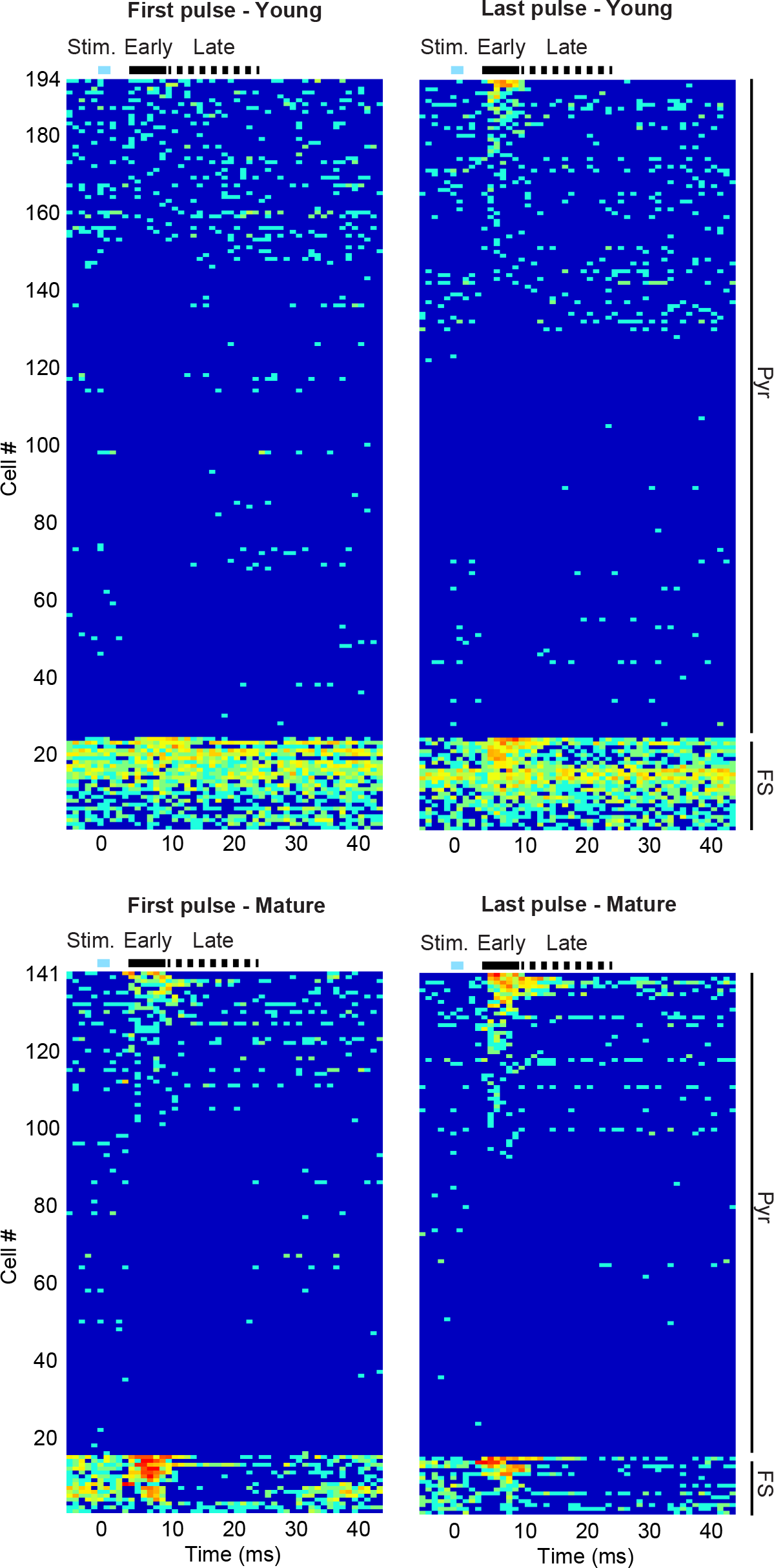
Mean responses evoked by the first or last train pulse. Activity (color coded) relative to aGC stimulation in the young (top; n=194) or mature (bottom; n=141) condition for all recorded cells (rows), ordered according to strength of response after the first (left) or last (right) train pulses and according to putative cell type (indicated). For young: Pyr=170, FS=24; for mature: Pyr=126, FS=15. Stimulation, early and late windows are indicated on the top left of the plot.

**Figure S11.**
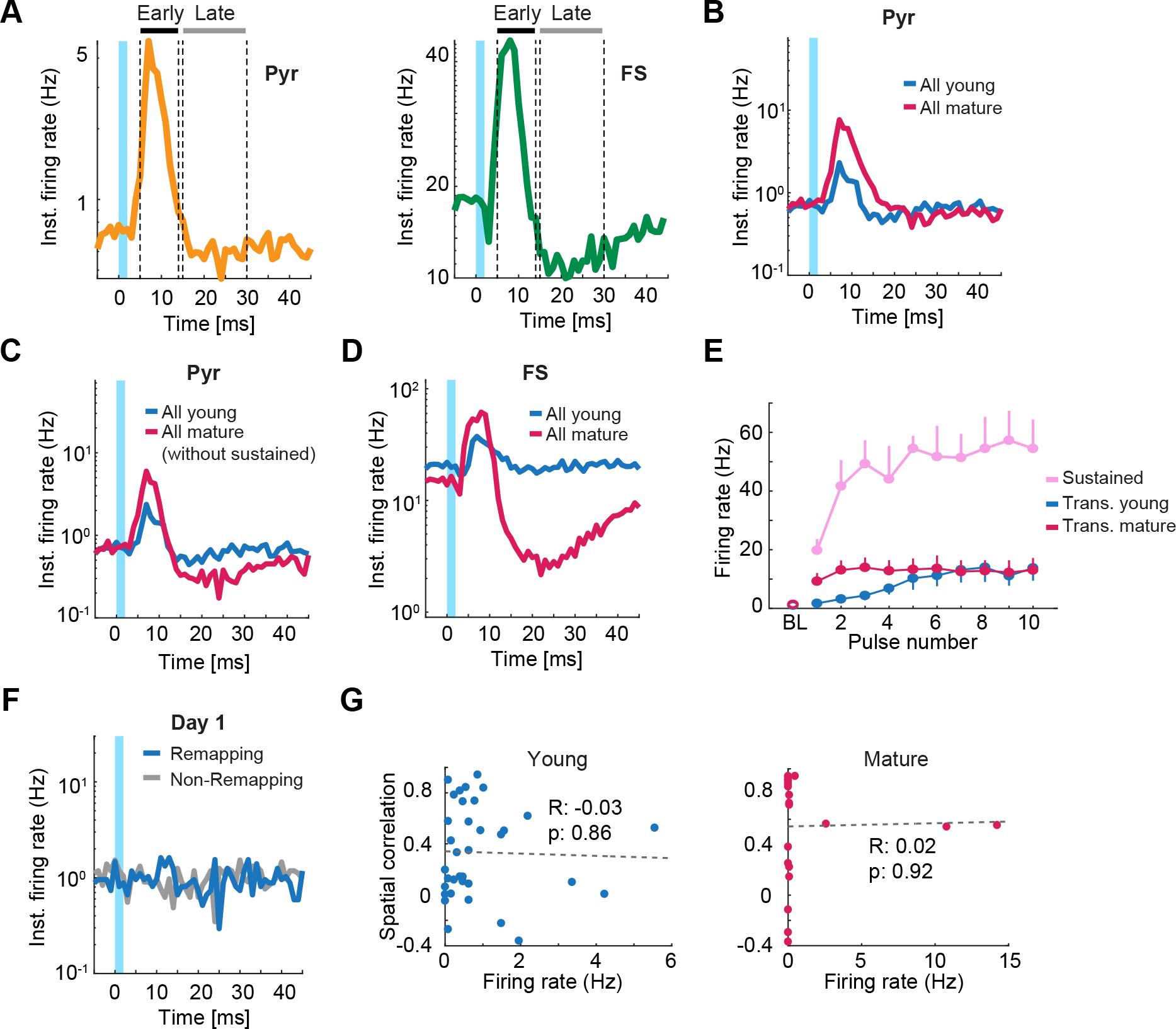
Patterns of CA3 activity evoked by young and mature aGC stimulation. (**A**) Mean evoked activity relative to optical stimulation onset (light-blue bar) for Pyr (left, orange; n=296) or FS (right, green; n=36) cells, after pooling data across days and conditions. Light stimulus (0-2 ms), early window (5-13 ms; black) and late window (15-30 ms; grey) indicated. Vertical dashed lines indicate the boundaries of early and late windows. (**B**) Mean evoked response over all recorded Pyr cells in the young (blue; n=170) or mature (red; n=126) conditions. Comparisons between conditions (shuffling): p = 0.0005 (early window), p = 0.23 (late window). (**C**) As B but excluding sustained activity cells (n=4). Note that differences between conditions in the early and late window are still observable. Comparisons between conditions (shuffling): p = 0.02 (early window), p < 0.0001 (late window). (**D**) Same as B but for FS cells (young=24; mature=15). Comparisons between conditions (shuffling): p = 0.03 (early window), p < 0.0001 (late window). (**E**) Evoked activity of transient young (blue), transient mature (red) and sustained (pink) Pyr cells along 10-pulse stimulation trains (mean ± s.e.m; BL: baseline). Comparison of ratio between last and first pulse vs. 1 for sustained cells (two-tailed Wilcoxon test): p = 0.125 (n=4). Note that, from the first to the last pulse, evoked response almost doubles. Other comparisons in Fig. 3D. (**F**) Mean evoked activity for remapping (blue, n=17) and non-remapping (grey; n=18) CA3 Pyr cells relative to young aGC stimulation on day 1. Comparisons between remapping and non-remapping cells (shuffling): p = 0.09 (early window), p = 0.58 (late window). (**G**) Spatial correlation (as in Fig. 1F) vs. late evoked activity on day 1 in the young (left; n=35) and mature (right; n=22) aGC conditions (regression coefficients indicated). In addition, we calculated the correlation (as in Fig. 1F) between remapping and the early evoked activity, as shown graphycally in this panel and in Figure 3E, but for the last pulse in the train. We found no statistical significance: young condition (r=-0.16; p=0.37), mature condition (r=0.03; p=0.88).

**Figure S12.**
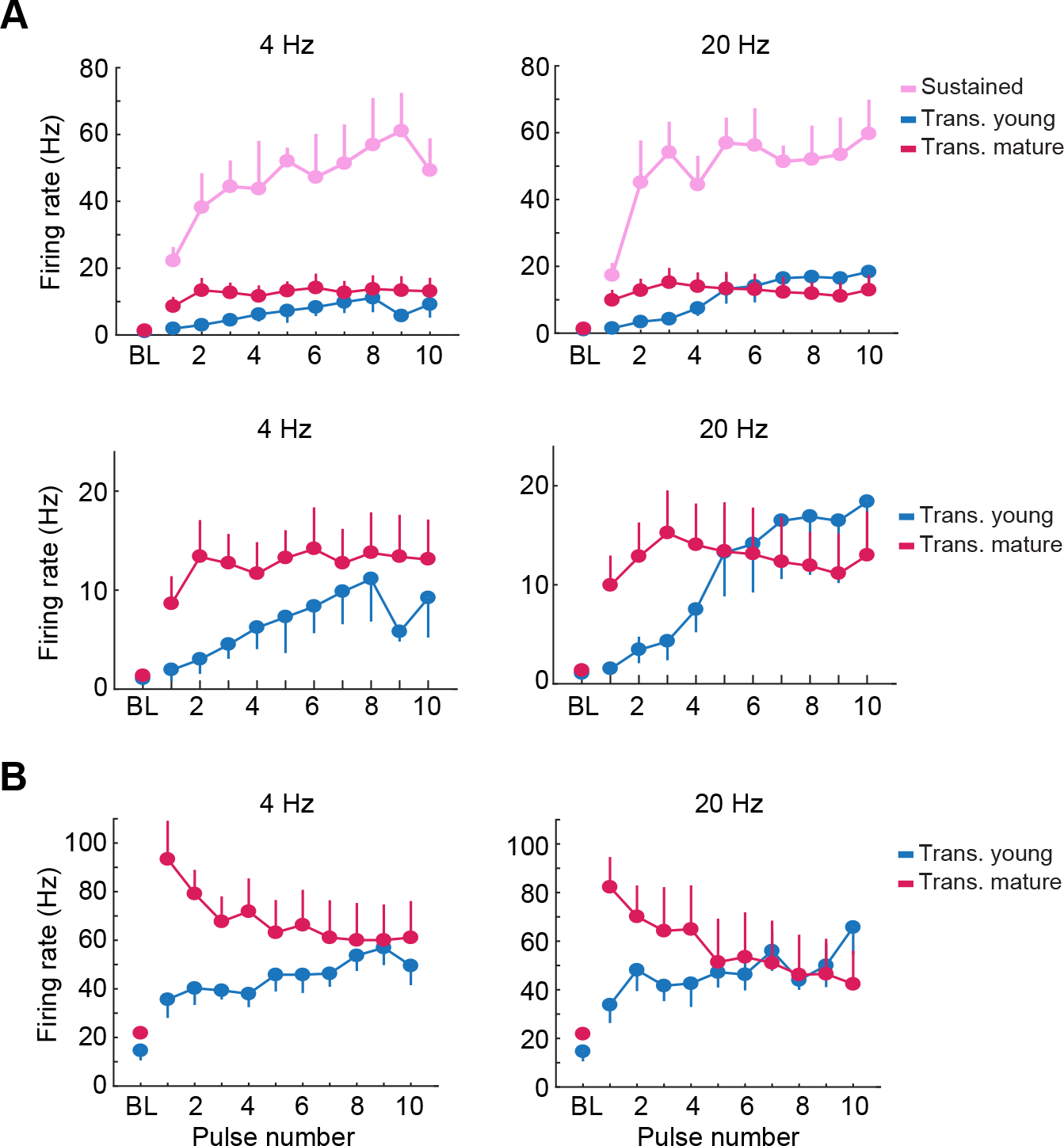
Similar short-term facilitation dynamic of responses evoked by 4 and 20 Hz stimulation trains. (**A**) Top: Early evoked activity of transient Pyr cells along 10-pulse stimulation trains at 4 Hz (left) and 20 Hz (right; mean ± s.e.m; BL: baseline; cells classified as in Figure 3). Comparison of ratio between last and first pulse vs. 1 (two-tailed Wilcoxon test): Pyr p = 0.03 (young; n=13), p = 0.006 (mature; n=21); FS p = 0.19 (young; n=7), p = 0.38 (mature; n=8). Sustained, p = 0.125 (n=4). Bottom: same but only for transient cells. (**B**) Early evoked activity of transient FS cells along 10-pulse stimulation trains at 4 Hz (left) and 20 Hz (right; mean ± s.e.m; BL: baseline; cells classified as in Figure 3). Comparison of ratio between last and first pulse vs. 1 (two-tailed Wilcoxon test): Pyr p = 0.0002 (young; n=13), p = 0.6 (mature; n=21); FS p = 0.16 (young; n=7), p = 0.2 (mature; n=8). Sustained, p = 0.125 (n=4).

**Figure S13.**
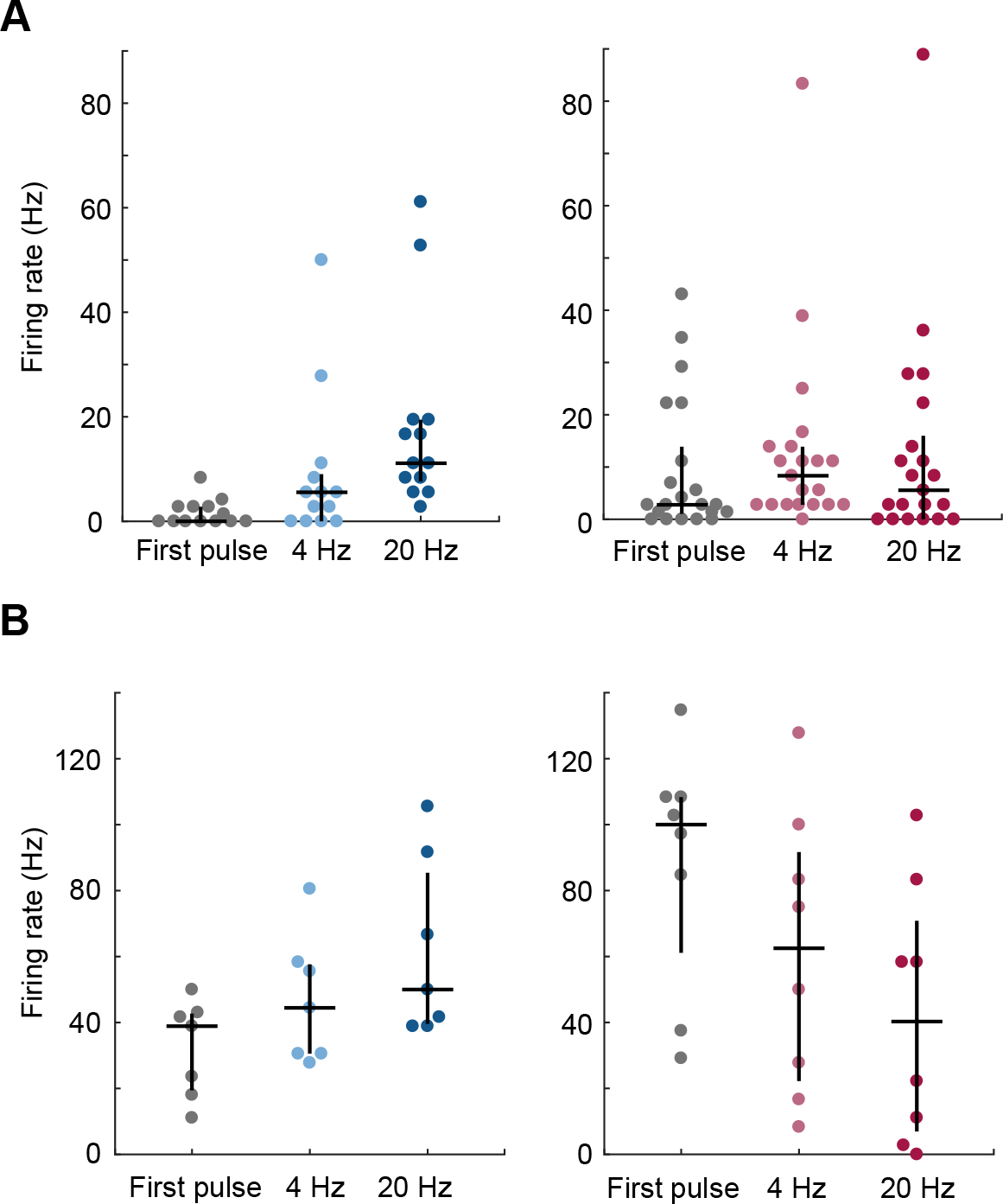
Frequency facilitation for Pyr cells in the young condition. (**A**) Evoked activity of transient Pyr across frequencies for young (left; n=13) and mature (right; n=21) conditions. From left to right in each panel: first pulse response pulled across frequencies (grey), last pulse responses at 4 (center) and 20 Hz (right). Friedman test: young χ2 (13) = 19.84, p = 0.00005, post hoc indicated that 20 Hz differed from 1 (p=0.00001) and 4 Hz (p=0.01); mature χ2 (21) = 2.46, p = 0.29. Results indicate a frequency facilitation effect in the young but not in the mature condition. Note that evoked response for the first pulse response in the mature condition is already high. (**B**) As in A but for FS for young (left; n=7) and mature (right; n=8) conditions. Friedman test: young χ2 (7) = 3.7, p = 0.16; mature χ2 (8) = 5.2, p = 0.07. Note, however, that responses tend to increase (young) or decrease (mature) with frequency. This agrees with reports of limited frequency facilitation below 20 Hz (Lee et al., 2019; Henze et al., 2002).

